# The 3D chromatin landscape of rhabdomyosarcoma

**DOI:** 10.1101/2022.12.05.519166

**Authors:** Meng Wang, Prethish Sreenivas, Benjamin D. Sunkel, Long Wang, Myron Ignatius, Benjamin Z. Stanton

**Author notes:** To whom correspondence should be addressed. (B.Z.S.), (M.I.). Joint Authors (Meng Wang, Prethish Sreenivas).

## Abstract

Rhabdomyosarcoma (RMS) is a pediatric soft tissue cancer with a lack of precision therapy option for patients. We hypothesized that with a general paucity of known mutations in RMS, chromatin structural driving mechanisms are essential for tumor proliferation. Thus, we carried out high-depth *in situ* Hi-C in representative cell lines and patient-derived xenografts to understand chromatin architecture in each major RMS subtype. We report a comprehensive 3D chromatin structural analysis and characterization of fusion-positive (FP-RMS) and fusion-negative rhabdomyosarcoma (FN-RMS). We have generated spike-in *in situ* Hi-C chromatin interaction maps for the most common FP-RMS and FN-RMS cell lines, and compared our data with patient derived xenograft (PDX) models. In our studies we uncover common and distinct structural elements in large Mb-scale chromatin compartments, tumor-essential genes within variable topologically associating domains, and unique patterns of structural variation. Our comprehensive analysis provides high-depth chromatin interactivity maps for contextualizing gene regulation events identification of functionally critical chromatin domains in RMS.

**HIGHLIGHTS:**

- PAX3-FOXO1 and MYOD localize in both A- and B-compartments
- Conserved mechanisms dictate CTCF orientation at TAD boundaries in RMS
- Differential TADs in each RMS subtype encompass tumor-specific genes
- Neo-TADs are formed from SV events in each subtype of RMS
- Both major RMS subtypes have structural variation that is identifiable from Hi-C
- Distinct mechanisms can produce the major fusion alleles in rhabdomyosarcoma
- PAX3-FOXO1 and MYOD genomic binding is more enriched at regions with CNV

## INTRODUCTION

Rhabdomyosarcoma (RMS) is an aggressive pediatric solid tumor, with prognoses that are highly dependent on the subtype (1-4). The major RMS subtypes are defined by histological features, termed alveolar RMS (ARMS), and embryonal RMS (ERMS), while more rare forms also exist (2,4). The ARMS subtype is often driven by oncoproteins encoded by fusion events resulting in *PAX3-FOXO1* or *PAX7-FOXO1* expression (5-7). ERMS can be driven aberrantly active signaling pathways, including RAS (8), HIPPO (9,10) and NOTCH (11) pathways, which play key roles. It is of note that the myogenic master regulator MYOD can drive RMS in its wild-type form (12-14), while mutations in MYOD drive a particularly aggressive and untreatable sub-type of RMS (2,15,16). Thus, understanding the molecular and chromatin-recognition functions of MYOD is of high interest for defining altered RMS epigenetics and disease etiology/tumorigenicity. Clinical studies have revealed that fusion-positive (FP)-RMS has poorer outcomes than fusion-negative (FN)-RMS (17,18). However, key questions remain. To what extent can chromatin architecture itself serve as a driver, to stabilize an aberrant epigenetic state? Are the genome structures of RMS subtypes conserved, despite the divergent clinical risk associated with fusion status? These key questions have motivated our comprehensive characterization and analysis of FN-RMS and FP-RMS, at the level of 3D chromatin structure and genome organization in cell lines and patient-derived tissues.

We sought to characterize genome structure-function relationships in RMS at several unique levels, including (**i**) larger (> Mb-scale) structural contact domains termed compartments (19-21), (**ii**) medium-range contact domains (< Mb-scale), or topologically associating domains (TADs) (19,22,23), (**iii**) in the context of the interactivity of copy number variation (CNV), and structural variations (SV) (24-27) and (**iv**) at the level of the gene regulatory loci encoding the major fusion oncogenes in FP-RMS (28). In our comparative analyses we discover substantial unique or subtype-specific regions at the compartment level (>1Mb) amid convergence in epigenetic conservation of FP-RMS and FN-RMS genomes. We report a comprehensive set of subtype-specific TADs, often bracketing tumor-associated (tumorigenic) genes. Within the *PAX3-FOXO1* locus in FP-RMS cells, we find evidence for unique structural events leading to the formation of the fusion alleles, which also has ramifications on local chromatin interactivity. Expanding upon this finding, in our structural analysis of the RMS landscape, we discover previously unknown SV events that are distinct even within and across RMS subtypes. We also uncover recurrent SV and CNV patterns within the ERMS subtype (29) in cell lines and patient derived xenografts (PDXs), suggesting that defects in genome integrity may be a common driver.

Our previous studies have focused on protein-centric chromatin conformation sequencing, modifying the HiChIP method in RMS for *in situ* spike-in (AQuA-HiChIP) of orthologous-species nuclei (14,30-32). Herein, we employed a modified version of AQuA-HiChIP allowing us to perform *in situ* Hi-C incorporating orthologous spike-in chromatin (Spice-C), to ensure that any absence of local chromatin interactions in our data was biological and not a product of experimental variation. Our 3D landscape presents a highly rigorous characterization of the initial high-resolution chromatin contact maps, and in-depth analysis of genome structure-function relationships in RMS. With our key definitions of conserved structural features in these tumors, we anticipate our studies will continue to catalyze impactful connections between primary genetic drivers and structural epigenetics.

## MATERIAL AND METHODS

### Cell Lines

The human Rhabdomyosarcoma cell lines RD (Female), Rh30 (Male), SMS-CTR (Male) and Rh4 (Female) were a gift from Dr. Peter Houghton (UTHSCSA), C2C12 Myoblasts (ATCC). All lines were maintained either in in DMEM or RPMI supplemented with 10% FBS at 37°C with 5% CO_2_. Cell lines were authenticated by genotyping.

### CRISPR/Cas9 knock out

Crispr guides at the PAX3 promoter were designed using Benchling® platform. The guides were cloned in pLentiCrisprV2-Puro or Hygro vectors. Lentivirus generated in 393T cells was used to infect RMS cells in the presence of 8μg/mL protamine and selected for 5 days in Puromycin (2μg/mL) + Hygromycin (100ng/mL) supplemented growth media.

### Cellular assays

Cell growth kinetics of *PAX3*-TAD KO cells were monitored using the Incucyte® imaging platform. Colony assays were performed by plating serially diluted control and KO cells in growth media and stained with Crystal violet after 15 days. Immunofluorescence staining was performed in cells after three days in differentiation medium, (DMEM+2% Horse serum) and fixed with 4% Paraformaldehyde, permeabilized in 0.5% Triton X-100/PBS and incubated with MyHC and MEF2C antibodies followed by Alexa-488/563 secondary antibodies along with DAPI and imaged using an Olympus FL-2000 confocal microscope at 20x magnification.

### Western blotting

Total cell lysate was obtained by lysing in 1x RIPA buffer supplemented with protease inhibitors (Roche) and MG132 (Calbiochem). Membranes (PVDF, BioRad) were developed using ECL reagent (Immobilon, Millipore). Internal control (GAPDH CST#2118) and target antibodies PAX3 R&D cat#MAB2457, FOXO1 CST cat#2880S, MYOD1 Santa Cruz cat#sc760 (western) CST #13812S (ChIP-seq), MYOG DSHB cat#F5D, MEF2C CST cat#5030, MyHC DSHB cat#MF20 were probed on different regions of the same blot and imaged (Li-Cor).

### ChIP-seq library preparation

Formaldehyde (1%, 10 minutes) fixed cells were sheared to achieve chromatin fragmented to a range of 200-700 bp using an Active Motif EpiShearSonicator. Chromatin samples were immunoprecipitated overnight at 4°C with antibodies targeting MYOD (CST cat#13812S), H3K27ac (Active Motif, cat#39133). DNA purifications were performed with modified ChIP protocol described previously^40^. For sample normalization 2 million C2C12 cells of were added to 6 million RMS cells before sonication. ChIP-seq libraries were prepared using Illumina TruSeqChIP Library Prep Kit. Libraries were multiplexed and sequenced using the NextSeq500 (Illumina).

### ChIP-Seq data processing

ChIP-seq data were processed with the ENCODE ChIP-seq pipeline with chip.xcor_exclusion_range_max set at 25. Specifically, Paired-end fastqs were aligned with bowtie2 to hg38, with parameters bowtie2 -X2000 --mm. Next, blacklisted region, unmapped, mate unmapped, not primary alignment, multi-mapped, low mapping quality (MAPQ<30), duplicate reads and PCR duplicates were removed. Peaks were called with MACS2, with parameters -p 1e-2 --nomodel -- shift 0 --extsize $[FRAGLEN] --keep-dup all -B –SPMR, where FRAGLEN is the estimated fragment length IDR analyses were performed on peaks from replicate samples or pseudo-replicates for MYOD ChIP-Seq with threshold 0.05. Motif analysis with HOMER(33) was then carried out on conservative IDR peaks. For visualization, bedGraph files were generated with MACS2 bdgcmp from the pile-up, and then converted to bigwig format with bedGraphToBigWig. Heatmaps were generated with deeptools. PAX3-FOXO1 ChIP-Seq were analyzed as previously reported.^40^ Raw sequencing data and processed files are available on GEO (GSE215203).

### Spike-in chromatin equalized Hi-C (SPICE-C)

RD, SMS-CTR, Rh30 and Rh41 cells were fixed for 10 minutes at room temperature (23 °C) and were spiked in with 20% mouse myoblasts (C2C12) and lysed gently with HiC buffer to release nuclei, permeabilized in 0.5% SDS for 10 minutes at 62 °C, quenched with 10% Triton X-100, and digested with MboI (200 U, 2h at 37 °C) which was then heat inactivated (20 min, 62 °C). Biotin incorporation was done with biotin-14-dATP (Thermo, cat# 19524-016) and DNA Polymerase I, Large (Klenow) Fragment (NEB, cat# M0210) for 1 hour at 37 °C. Then we performed *in situ* ligation with T4 DNA ligase (4h room temperature). Nuclei were resuspended in TE and sonicated (5 cycles with shearing ‘on’ time with 30 seconds ‘on’ 30 seconds off), using the Active Motif Epi-shear probe sonicator at 30% power. The sheared DNA was reverse crosslinked with SDS and Proteinase K O/N and pulled down with M-280 Streptavidin Dynabeads (Thermo, cat#11205D), washed, eluted followed by end repair, A-tailing, adapter ligation and library amplification all on-bead as previously reported ^7^. For a detailed protocol of the SPICE-C method refer to **Supplemental Methods** section.

### Hi-C Sequencing

Libraries were indexed with Illumina barcodes and quantified using *Qubit* (ThermoFisher), with concentrations measured between 1-10ng/μL. Median fragment lengths for our libraries were measured with *TapeStation* (Agilent) and were between 200bp-500bp with average sizes approximately 350 bp for each indexed library. After validating indexed library concentrations and indexes with qPCR (IGM, Nationwide Children’s Hospital), we sequenced the libraries on the *NovaSeq – S1* platform (Illumina) with paired-end 150 bp reads and 300 sequencing cycles (∼1 billion reads) and demultiplexed for further analysis. Two independent biological replicates for each cell line were combined informatically to get contact maps with 30 million valid, long-range cis contacts.

### Hi-C data processing

Hi-C data are processed with accordance to 4D Nucleosome processing pipeline recommendations. Specifically, fastq-files are aligned to hg38 with bwa mem. Mapped reads are then tagged for PCR duplications, parsed for ligation junction, merged, and then sorted into binned matrix using pairtools. Matrix aggregation and normalization (ICE) were carried out with cooltools.

### Hi-C data analyses

Compartments are called at 100kb using cooltools call-compartments, using H3K27ac as reference track. TADs are called with R package TopDom. Differential TAD analysis is carried out with R package diffHic. We used hicbreakfinder to find structural break points, and then neoloopfinder to estimate copy number and reconstruct structural variations. We used hicexplorer, juicebox and HiGlass for Hi-C data visualization.

### Versions of Software used

Versions of Software used:
|  |  |
| --- | --- |
| **bwa** | **0.7.17-r1188** |
| **pairtools** | v0.3.0 |
| **cooltools** | v0.4.0 |
| **topdom** | v0.10.1 |
| **hicexplorer** | v3.7 |
| **neoloopfinder** | 0.2.4.post2 |
| **R** | v4.0.0 |
| **juicebox** | 2.2.2 |
| **diffHic** | v1.22.0 |
| **hicbreakfinder** | NA |
| **ENCODE ChIP-Seq pipeline** | v2.1.2 |
| **deeptools** | v3.3.1 |
| **HOMER** | v4.11.1 |

## RESULTS

### Self-interacting domains in rhabdomyosarcoma

In our previous studies, we developed an orthologous chromatin spike-in approach for short-range chromatin interaction domains (31). However, the resolution in these studies precluded our ability to define large scale compartmental structures (19,20,34) and smaller scale TADs. We were motivated to understand chromatin domains in rhabdomyosarcoma, and thus established our approach to perform *in situ* Hi-C for deep sequencing. With the same experimental workflow as in our previous studies (14,31), we developed a modified *in situ* Hi-C protocol Spice-C (see **Methods**) (19). In our approach, we permeabilize nuclei with gentle SDS treatment and heat (32), perform restriction digests with MboI, biotinylate exposed DNA ends with a Klenow extension, and perform proximity ligation to stitch together proximal chromatin ends associated with *cis*-interactions. We gently shear chromatin prior to reverse crosslinking and on-bead library preparation for next-generation sequencing. We noted that our early attempts to scale up the *in situ* Hi-C reactions resulted in quenching of the library amplification step with our streptavidin beads, which is a common caveat of on-bead library preparation. We circumvented this issue through preparation of individual amplification reactions with no more than 10 μL beads in sequencing library amplifications (see **Methods**).

We established our modified *in situ* Hi-C method for FP-RMS cells (Rh4, and Rh30) and FN-RMS cells (RD, and SMS-CTR) with approximately 1 billion paired-end sequencing reads per cell line after combining replicate samples. We mapped the reads onto the human reference genome, tagged for PCR duplication removal, and parsed into contact pairs. We binned valid contacts into interaction matrices and normalized interaction matrices using the ICE algorithm (35) for subsequent analyses.

### Analyses of Compartment Structures in RMS

Compartments are megabases (Mb) long, self-interacting large scale chromatin structures (20). Generally, human genomes can be classified into “compartment A” and “compartment B” via principal component analysis (PCA) of the interaction matrix (19-21). Regions in compartment A are gene-rich and are usually in an “active” local epigenetic state. These regions usually exhibit higher GC-content and contain more active chromatin markers such as H3K27ac (21,36,37). In contrast, regions in compartment B are relatively gene-poor and show higher levels of silent or repressive local epigenetic markers such as H3K9me3 (38,39). Based on these previous findings (19,20), we carried out compartment analysis with our *in situ* Hi-C at 100 kb resolution. Compartments were assigned based on the sign of the first principal component (PC1) value of the correlation matrices, using H3K27ac as the reference track (19,20). We generated compartment definitions for two fusion-driven RMS cell lines (Rh4, Rh30; **Figure 1a,b**), and two fusion-negative cell lines (RD, SMS-CTR; **Figure 1c,d**) and developed these maps for comparative analyses.

**Figure 1.**
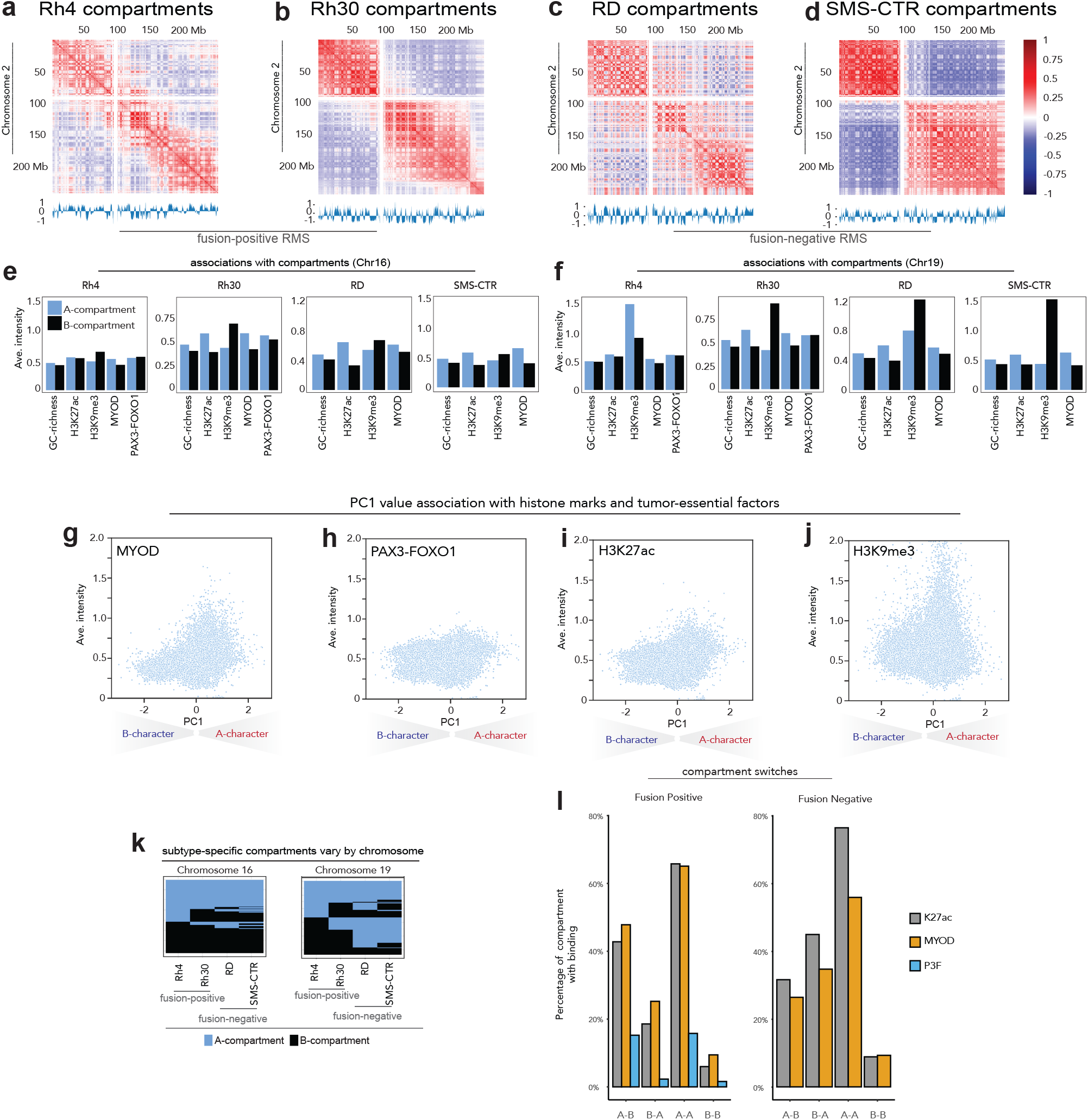
Chromatin compartments in Rhabdomyosarcoma. Correlation heatmaps along chromosome 2 in (**a**) Rh4 cells, (**b**) Rh30 cells, (**c**) RD cells, and (**d**) SMS-CTR cells, with normalized eigenvectors shown below, and correlation values shown at right. The associations between compartment calls and GC-richness, H3K27ac, H3K9me3, MYOD binding, and PAX3-FOXO1 binding are shown for Rh4, Rh30, RD, and SMS-CTR cells on (**e**) chromosome 16 and (**f**) chromosome 19 with A-compartments shown in grey and B-compartment shown in orange. PC1 values for compartments are plotted against average intensity for chromatin marks and regulatory factors: H3K27ac (**g**) H3K9me3 (**h**), MYOD as scatter (**i**) and PAX3-FOXO1(**j**). (**k**) A-compartments and B-compartments are plotted in blue and black, respectively for Rh4, Rh30, RD, and SMS-CTR on chromosome 16 (left) and chromosome 19 (right). (**l**) Discordant and Concordant compartments’ association with deposition of H3K27ac, MYOD and PAX3-FOXO1. Discordant compartments were defined as A-B (Compartment A in FP and Compartment B in FN) and B-A (Compartment B in FP and Compartment A in FN).

To understand the roles of chromatin modifications in the compartmental structures of FP-RMS and FN-RMS, we established H3K27ac, H3K9me3 ChIP-seq in each cell model, using gentle sonication to avoid epitope shearing (40,41). We sought to provide mechanistic linkage between large scale architectural features and modifications on the linear epigenome that may be intrinsic to subtype. We found that regions belonging to compartment A have a higher level of GC content compared to regions assigned to compartment B (**Figure 1e,f**) with exceptions within chromosomes 3, 12 and 18 (see **Supplementary Figure S1-S4**). However, H3K9me3, a repressive histone modification, is not always enriched in compartment B across chromosomes (**Figure 1e,f**; **Supplementary Figure S1-S4**). While the intensity of H3K9me3 is systematically higher in compartment B across many chromosomes, this histone mark is also present in compartment A, especially noticeable at chromosomes 5, 18, 21, and others (**Supplementary Figure S1-S4)**. To our surprise, we also observed substantial signal of H3K27ac and H3K9me3 in both compartmental classes. This is consistent with histone modifications having unique roles in a context-dependent fashion, or with heterochromatin regions being punctuated with activating marks as has been shown by us and observed in other cell types (37,42). The relative enrichments of H3K9me3 in B-compartments were more pronounced in the fusion-negative subtype (**Figure 1e,f**, **Supplementary Figure S1-S4**). The greater degree of concordance between deposition of H3K9me3 and B-compartments in FN-RMS relative to FP-RMS may be related to distinct large-scale compartment structure in each subtype, or alternative functions of H3K9me3 and heterochromatin formation in a subtype-specific context. As we used H3K27ac to assign compartmental identities, it follows that some chromosomes unexpectedly exhibit a positive correlation between levels of H3K27ac and H3K9me3. This unexpected concordance may have resulted from locally distinct epigenetic functions of H3K9me3, which can reside both within active and repressive chromatin regions (41,43). We noted that while GC-richness is invariant in the context of local or global epigenetic state, deposition of histone marks is highly dependent upon the local chromatin context (**Supplementary Figure S1-S4;** (35,44).

*PAX3-FOXO1* is a defining tumor driver for fusion-positive RMS (6,7), and the *MYOD1* gene product has also been shown to be among the most essential in FP-RMS (14). Mutations in *MYOD1* result in an incurable form of the disease (2,16). To understand chromatin recognition preferences and to assess if there exists a high degree of association between chromatin compartmental domains and these factors, we established genome-wide ChIP-seq profiles for each of these major tumor drivers, using our gentle sonication approach (41). Since PAX3-FOXO1 has been shown to have pioneer transcription factor function, and MYOD may have pioneer factor-like properties that augment its association with compacted chromatin regions (**Supplementary Figures S1-S4**), (40,41) we specifically asked whether MYOD and PAX3-FOXO1 would also be highly associated with compartment B in RMS (14,31). We observed PAX3-FOXO1 and MYOD within both compartments A and B (**Figure 1e,f**; **Supplementary Figures S1-4**), although PAX3-FOXO1 is more associated with B compartment relative to MYOD (**Figure 1g,h**). These are novel findings showing that these regulatory factors can associate with both open and repressive chromatin domains (30).

We next asked whether compartment strength was associated with RMS subtype. We analyzed the compartmental distribution data and generated saddle plots (35,45,46), which revealed intra-compartmental and inter-compartmental interaction preferences in RMS cell lines. In all four RMS cell lines there exists a preference for self-interaction within compartment class (e.g., B-B interactions or A-A interactions), relative to cross-compartmental interactions (**Supplementary Figure S5**). Moreover, we found that for each RMS cell line, regardless of fusion status, B-B compartmental interactions were stronger than A-A homotypic interaction strength. While these trends suggested epigenetic convergence across FP- and FN-RMS, we did observe that Rh30 had a higher prevalence for inter-compartmental interactions across compartment classes, relative to the other RMS cell lines that we investigated (**Supplementary Figure S5**). The unique compartmental architecture in Rh30, with increased tendency for chromatin interactions occurring across compartment classes, suggests that while there is a high degree of subtype-specific structure, variation also exists within subtypes.

### Association with Transcription Factors and Histone Marks

We next sought to understand the concordance between PC1 (35,46) and MYOD, PAX3-FOXO1, H3K27ac, and H3K9me3 signals. We observed that the sign and magnitude of PC1 is indictive of levels of H3K27ac and H3K9me3 depositions, in opposite directions (**Figure 1i-j; Supplementary Figure S6**). The larger positive PC1 values were associated with higher levels of H3K27ac signal, whereas H3K9me3 signal does not exhibit the same association (**Figure 1i,j**). The general trend of increased signal level as a function of increased magnitude of PC1 values was observed for MYOD in compartment A, generally more reminiscent of H3K27ac (**Figure 1g,i**). However, PAX3-FOXO1 had relatively consistent signal irrespective of the sign or magnitude of PC1 except for the strongest A-compartments which exhibited increased binding (**Figure 1h**). These data are again consistent with an ability of PAX3-FOXO1 to bind both active and inactive chromatin across a wide variety of chromatin structural domains (40,41). These data suggest that the master regulatory factors PAX3-FOXO1 and MYOD have distinct chromatin structural roles in RMS, as opposed to overlapping functions.

### Identification of subtype-specific compartments

We hypothesized that compartments that switched classification between FP and FN cell lines would be salient when comparing across RMS subtypes. To investigate the similar (concordant) and dissimilar (discordant) compartments in each major subtype, we defined the subtype-specific compartments in FP and FN RMS cell lines. Overall, we found that over 83% of the genome in each RMS cell model has been assigned a compartment, and 55% of all compartments are consistent in all four FP and FN cell lines. 72% of FP-RMS compartments were consistent within the two cell lines (Rh4, Rh30), while FN-RMS cell lines (RD, SMS-CTR) showed a higher level of consistency at 82% (**Figure 1k, Supplementary Figure S7**). In terms of the subtype-specific compartments, Concordant A compartments (A-A) constituted 30.0% of all compartments, while concordant B compartments (B-B) accounted for another 25.0% of all compartments. Discordant compartments were defined as A-B (Compartment A in FP and Compartment B in FN) and B-A (Compartment B in FP and Compartment A in FN). Even though discordant compartments constituted a relatively small proportion of all compartments (6%), we did observe some stereotypic patterns of compartment discordance. Subtype-specific compartmentalization was especially prevalent on chromosomes 19 and 22 (17.9% and 17.4% respectively), while chromosomes 17, 21 and 1 had more compartmental similarity across subtypes (74%, 73.2% and 72.4% respectively; **Figure 1k, Supplementary Figure S7**, *blue = compartment A, black = compartment B*).

We next proceeded to examine if chromatin regulatory factors were associated with discordant compartments. We examined the prevalence of histone acetylation at A-B, B-A, A-A and B-B regions. We found that H3K27ac has the highest concentration in A-A compartments. 65.8% of FP and 76.4% of FN A-A regions have at least one H3K27ac site. In comparison, only 5.98% of FP, and 8.86% of FN B-B regions have H3K27ac **(Figure 1l**). The stark contrast suggests that regions with consistent compartment assignments might share similar structures and functions between subtypes. In discordant compartments, H3K27ac deposition is higher in the A compartment which is consistent with the definition. Overall, MYOD shows a similar profile as H3K27ac, both in percentage of compartments occupied and the general trend of preferential binding within A compartments. This is consistent with MYOD signal intensity (**Figure 1g**) tracking closely to H3K27ac (**Figure 1i**). However, MYOD does occupy more B-B compartments and fewer A-A compartments than H3K27ac. Moreover, we find that at MYOD sites associated with B-to-A switching from FN-to-FP are associated with Wnt signaling molecules (**Supplementary Table S1**). PAX3-FOXO1 is less commonly observed in A-A and A-B compartments, mostly because PAX3-FOXO1 has much fewer FP-consensus binding events throughout the genome. While H3K27ac and MYOD are more common in A-A compartments, PAX3-FOXO1’s presence in both compartments is almost equal, showing a higher affinity to regions that changed from B to A. This is an extension of our findings that PAX3-FOXO1 has pioneer function in FP-RMS cells (40,41).

### Analysis of topologically associating domains (TADs) in RMS

Human genomes contain structures that are highly self-interacting and self-insulating (22,23). These structures, initially observed by Heard and Ren and co-workers, span tens of thousands of base pairs to a few Mb, and are known as Topologically Associating Domains (TADs). To form TADs, CTCF and cohesin function together via a loop extrusion mechanism (35), with cohesin facilitating the loop extrusion and CTCF functioning as an insulator to block cohesin processivity (19,34,47,48). Interestingly, when CTCF is conditionally deleted, TADs are lost, while large scale compartments persist, suggesting a decoupling of shorter range and longer-range chromatin organization (47). There is evidence that TADs are generally conserved across mouse to human chromatin, and conserved across tissues within human cells lines (19). However, we hypothesized that identification of subtype-specific TADs for RMS would allow us to conceptually and mechanistically connect chromatin structure and gene expression regulation in the tumor. Thus, we studied the chromatin domain structure of FP and FN-RMS, with a focus on the differential TADs between the subtypes.

We observed that TAD assignments are very sensitive to many parameters, among which include resolution of bins and size of the moving window. For our analysis, we used 50kb bins and a moving window of 10-fold on each side, resulting in median TAD size of approximately 1Mb, which is consistent visually (**Figure 2a,b**). We uncovered 1860, 1920, 2019 and 2059 TADs for Rh4, Rh30, RD and CTR cell lines, respectively. Most of the TADs are consistent within all four cell lines and have CTCF binding events at both TAD boundaries. For example, the TAD at chr2: 216.75Mb – 217.75Mb is present in all four RMS cell lines, demarcated with strong CTCF signal (**Figure 2c**). Even though the strengths of the TADs differ slightly across cell lines, we observe the signature “dot structures” formed by the two TAD-defining CTCF binding sites at the boundaries in all four RMS cell lines (34,47,49). We also observe the distinct stripe pattern which may be evidence of cohesin extrusion in action (50). The sub-TAD strengths of CTCF-demarcated loop domains are relatively weaker in the RD cell line than in others, but sub-TAD boundaries at around 217.20Mb and 217.50Mb are both visible, as exemplified in a highly representative TAD structure (**Figure 2c**).

**Figure 2.**
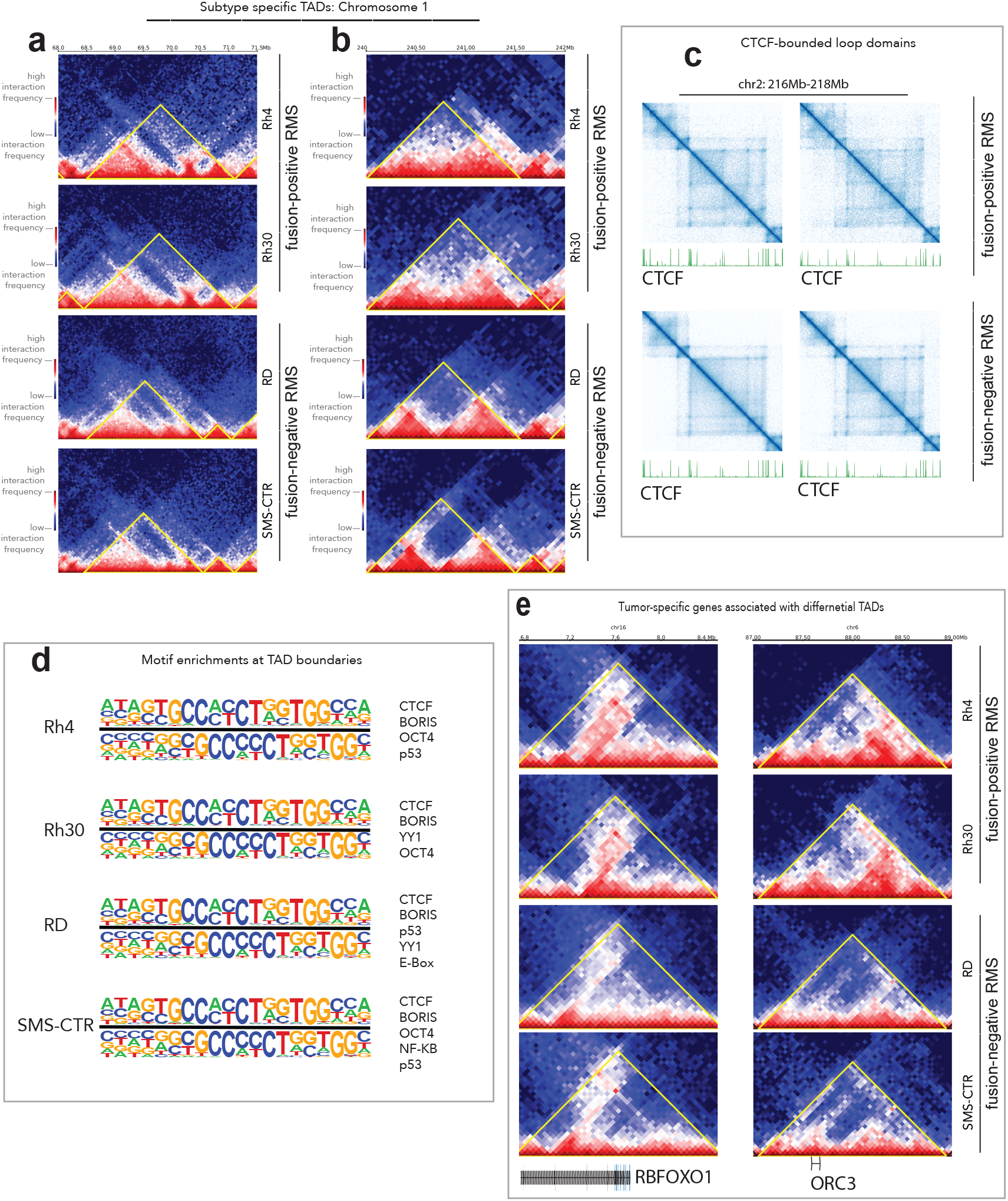
Topologically associated domain in Rhabdomyosarcoma. (**a, b**) Subtype-specific TADs are shown on chromosome 1 for Rh4, Rh30, RD, and SMS-CTR with a colorimetric interaction intensity scale (shown, left) and subtypes (shown, right). (**c**) CTCF-bounded loop domains are shown with interaction intensity plotted in blue above CTCF ChIP-seq data for each cell line in green. (**d**) Motif enrichments at TAD boundaries are indicated for Rh4, Rh30, RD, and SMS-CTR. (**e**) Tumor-specific genes associated with differential TADs in our RMS cell models are indicated.

We sought to determine if there exists association between PAX3-FOXO1 and the genomic loci encoding boundaries. Based on recent studies in the muscle lineage (51) illustrating MYOD localization at loop anchors, we hypothesized that MYOD, a critical core regulatory factor essential for myogenic cell fate determination, would be localized to TAD boundaries. We found only moderate associations of the canonical E-Box motif localized to these boundaries and there was no evidence of MYOD preferentially binding to TAD boundaries (permutation test; *p*-value <= 0.67). To our surprise, there was no evidence that PAX3-FOXO1 binding events were significantly enriched at the TAD boundaries (permutation test; *p*-value <= 0.99). The most highly enriched motif associated with TAD boundaries was CTCF (adjusted *p*-value < 10^−4^). Both P53 and YY1 motifs were enriched within our datasets but not significantly (**Figure 2d**). Thus, we find evidence for CTCF as instructive for TAD maintenance in FN-RMS and FP-RMS, as well as the conservation of key motifs, including YY1 for TAD domain boundaries across subtypes. We attribute these findings to a fundamental distinction between loops and TADs. While some loop domains are synonymous with TADs, many can be differently defined based on the fundamental properties of the class of *cis*-chromatin interaction (19,34). In comparison, CTCF binding sites are highly concentrated at TAD boundaries (permutation test, *p*-value < 10^−5^; **Figure 2d**). These results are in accordance with previous reports in other tissues where associations between CTCF/Cohesin were found at boundaries (19,34,50,52). Understanding mechanisms of PAX3-FOXO1 and MYOD for shorter-range looping events will be of high interest in future studies. Mechanistically, it is interesting that the functions of these essential tumor driver TFs may exist at the larger compartment level, while being more context-specific or gene-specific for TAD structures.

To investigate if TADs are associated with gene expression patterns in RMS, we analyzed 303 differentially called TADs between FP and FN RMS cell lines (FDR < 0.01, FC > | log_2_(1.5)|). Some TADs that display distinct patterns between subtypes are located close to genes associated with tumorigenesis, such as *RBFOXO1, ORC3, TCF4, PTN and NRG1* (**Figure 2e; Supplementary Figure S8**). We observed *RBFOXO1* showing stronger gene body and downstream interactions in FP than in FN RMS cell lines. For *ORC3*, there exist stronger interactions between upstream regions and the gene body in FP than in FN RMS cell lines. In our compartment analysis we also observed that the genes that are involved in A-B or B-A switching mostly encompassed a neurogenesis signature (128 genes) by GSEA analysis, while a small subset included genes involved in myogenesis (11 genes, **Supplementary Table S2**). Importantly, a majority of differential TADs between FP and FN-RMS were associated with genes involved in neurogenesis. Our findings provide a unique chromatin structural context for studies that find that FP-RMS tumors can express genes involved in neurogenesis (53,54).

### Genome integrity and RMS: CNV, SV, and classical fusions

Structural variations (SV) include genomic alterations such as somatic mutations, genome duplication, genome deletion and genomic translocation events. Recent studies have revealed that SVs and chromosomal imbalances are prevalent in cancer (25,55,56), and specifically in childhood tumors (57-59). For example, there is evidence that SVs occur in diffuse intrinsic pontine glioma (DIPG), leukemias, lymphomas, and is relatively enriched in childhood sarcomas (29,57-62). In RMS, certain CNVs, including *MYCN* amplifications (2), have been associated with chemotherapy resistance and poor patient prognoses. Interestingly, it has been reported that FN-RMS genomes go through more structural alterations compared to FP-FMS genomes, while FP-RMS often undergoes genome duplication events in patients (27,29).

We applied *neoloopfinder* (63) to reconstruct copy number profiles for our FP (Rh4, Rh30) and FN (RD, SMS-CTR) RMS cell lines. We found pervasive copy number changes in all four RMS cell lines regardless of fusion status (**Figure 3a-e**; **Supplementary Figure S9**). The Rh30 genome has the highest percentage of altered copy number, as only 46.51% of the Rh30 genome remained copy number neutral. The SMS-CTR cell line has the next highest level of altered copy number, with 48.31% of the genome remaining copy number neutral. Rh4 and RD cell lines are slightly more stable, with 54.86% and 55.42% of the genome being copy number neutral, respectively. At the same time, Rh30 has slightly more single copy loss than the other cell lines, with 30.54% of the genome estimated to have CN=1 (**Figure 3f-m**). Strikingly, PAX3-FOXO1 and MYOD both localize to CN-enriched loci in Rh4 cells, suggesting that primary and essential tumor drivers can interact with neo-scaffolding elements occurring through genome instability (**Figure 3l,m**). It is of note that CNV in RMS is associated with the gene loci encoding *PAX3-FOXO1* and also provide scaffolding for PAX3-FOXO1 binding (**Figure 3f,g,m**, **Supplementary Table S3**). We hypothesize that this reveals a mechanism where CNV both enhances tumor-essential gene dosage and also provides scaffolding for the gene products to localize. Also, of note is that *MYCN* copy number is elevated in Rh30 cells, potentially connecting high MYCN expression with structural alterations (**Figure 3g, Supplementary Table S3**). We observed striking elevations in CNV at the loci encoding the cell cycle regulator *CDK4* (**Supplementary Table S3**), which is associated with more severe patient outcomes in FP-RMS (2). Even though copy number profiles differ from cell line to cell line, there exist deletions that are common to both FN and FP cell lines, such as the deletion at the start of chr22 (**Figure 3e**) and amplification at the start of chr14 (**Supplementary Figure S9**). Our work has revealed key patterns of SV and CNV in each major subtype, which provides new evidence that events leading up to but not limited to the canonical fusion events in RMS may be critical for sarcomagenesis. We anticipate that CNV calls from our deep 3D genomic sequencing will continue to serve key roles in connecting chromatin architectural organization and structural alterations in human cancer.

**Figure 3.**
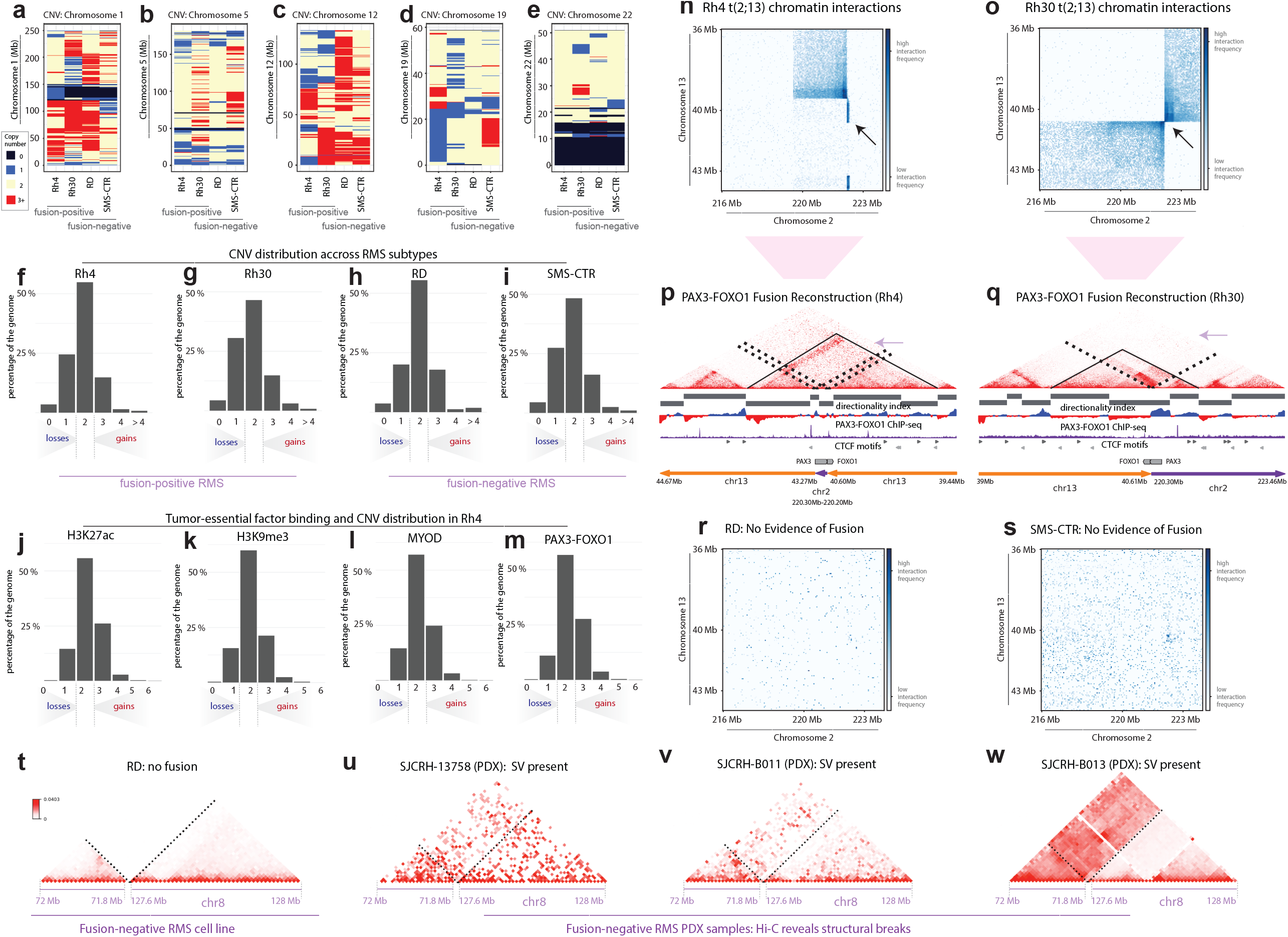
Copy number and structural variations from Hi-C. Copy number variation (CNV) is indicated with colorimetric scale for Rh4, Rh30, RD, SMS-CTR for chromosome 1 (**a**), chromosome 5 (**b**), chromosome 12 (**c**), chromosome 19 (**d**), and chromosome 22 (**e**). On the scale, black = deletion/loss, blue = single copy, beige = two copies, and red = 3 or more copies. The distributions of CNV gains and losses are shown for Rh4 (**f**), Rh30 (**g**), RD (**h**), and SMS-CTR (**i**) as a function of percent coverage across the genome on the y-axis. Fusion status for each cell line is indicated beneath the plot. The association of chromatin regulatory factors and histone marks with CNV in Rh4 cells is shown as a function of disposition of H3K27ac (**j**), H3K9me3 (**k**), MYOD binding (**l**), or PAX3-FOXO1 binding (**m**). Heatmaps detailing the translocation at PAX3 (chr2) and FOXO1 (chr13) loci in Rh4 (**n**) and Rh30 (**o**) FP cell lines, as well as background interactions at RD (**r**) and SMS-CTR(**s**) FN cell lines. Reconstruction of PAX3-FOXO1fusion gene locus and neo-TADs in Rh4 (**p**) and Rh30 (**q**) cell lines. Recurrent patterns of structural variation along chromosome 8 are not present in FN-RMS cells, RD (**r**), while they are present with increased interaction frequencies between 71.8 Mb – 127.6 Mb for FN-RMS PDXs, SJCRRH-13758 (**s**), SJCRRH-B011 (**t**), and SJCRRH-B013 (**u**).

We next asked whether SV identification from Hi-C could illuminate structural attributes in the locus encoding the major tumor driver in FP-RMS. Another type of structural variation is genome reconstruction resulting from a break in the linear chromosome fiber. A failure to repair the break could result in a translocation event, a deletion, or an inversion (reviewed(25)). In FP-RMS, SVs lead to fusion genes such as *PAX3-FOXO1* and *PAX7-FOXO1*, which are essential in FP-RMS but poorly understood at the molecular level. We asked the questions of whether (**i**) the *PAX3-FOXO1* fusion locus in Rh4 and Rh30 cell lines is structurally the same, and (**ii**) if there are structural breaks common to FN cell lines. Using bioinformatic tools, we were able to reconstruct and visualize *PAX3-FOXO1* fusion locus along the altered genome (24,63). As one would expect, the chromosomal break is visible in Rh4 and Rh30 cell lines but absent in RD and SMS-CTR cell lines (**Figure 3n, o, r and s**). The SV event in Rh30 cell line is a translocation between TTS of *PAX3* and TSS of *FOXO1*. This is consistent with the interaction heatmap showing a symmetric radiating pattern originates at chr2:222.2Mb and chr13:40.6MB (**Figure 3o,q**). The SV event resulting in *PAX3-FOXO1* in the Rh4 cell line, however, is more complex. The SV involves an insertion of the *PAX3* gene (chr2:222.2Mb-222.3Mb, anti-sense) inserted between chr13:40.6Mb and chr13:43.27mb, producing the two strip-like hotspots on the heatmap (**Figure 3n**). However, the square to the top left of the break site also suggests an inversion at chr2:222.2mb, just downstream of *PAX3*, and chr13:39.5Mb. Furthermore, there also exist long range interactions of chr3 between 41.25Mb and 77.00Mb, as well as inter-chromosomal interaction between chr2:222.0Mb and chr13:77.50Mb **(Figure 3p, Supplementary Figure S10)**. The interactive architecture of chr2 and chr13 in Rh4 cell line is not well defined, as multiple SVs could exist from independent fusion or amplification events.

We also studied TADs on the reconstructed Rh4 and Rh30 genomes (neo-TADs) by calculating directionality index (DI; (22)) on the reconstructed genome. In both Rh4 and Rh30 cell lines, the *PAX3-FOXO1* fusion gene is located at the inside of a neo-TAD (**Figure 3p,q**). In the Rh4 cell line, the TAD spans chr13:39.6Mb-40.6Mb, chr2:222.2Mb-222.3Mb, and then chr13:43.27Mb-43.9Mb, with CTCF binding sites flanking both ends of the TAD and pointing to the interior of the TAD. We have discovered that convergent CTCF sites flanking the *PAX3-FOXO1* neo-TAD are in agreement with the high degree of CTCF-motif convergence flanking TADs across mammalian chromatin architectural systems (19). In the Rh30 cell line, the neo-TAD spans 39.6Mb to 40.6Mb on chr13, and then 222.2Mb to 222.7Mb, with evidence suggesting a sub-TAD from chr13:40.0Mb-chr2:222.7Mb (**Figure 3q**). Thus, despite the common genetic drivers in FP-RMS, the events leading to the SV encoding the fusion alleles has occurred through distinct rearrangement mechanisms. We extended our analyses of SV into patient-derived xenografts (PDXs) from the FN-RMS subtype to understand the context for frequent SV events reported in this subtype (29). We investigated the structural variations in PDXs from FN-RMS patients obtained from St Jude Research Hospital (SJCRH-13758, SJCRH-B011, and SJCRH-B013). Using the same algorithms for SV detection, we found a structural break located on chr8 in all 3 PDX samples, near the *MSC* gene. while the same break is not conserved in RD or SMS-CTR cell lines (**Figure 3t-w**). Thus, high-depth Hi-C in RMS has revealed unique patterns of SV in each cell line or PDX, while some of these SV events produce convergent gene fusions, and others produce divergent structural and architectural features.

We identified chromatin contact domain structures near the *PAX3* locus which we hypothesized are essential to our FP RMS cell lines (**Figure 3n,o; Figure 4a**). *PAX3* is an important regulator of stemness in the muscle (64,65). In fusion-positive RMS the *PAX3-FOXO1* oncogene is a driver of this disease (6,7), while a role for *PAX3* in fusion negative RMS remains to be defined. We hypothesized that these conserved domain structures had intrinsic functional significance for RMS tumor maintenance. We noted that there are distinct contact domains at the *PAX3* locus in Rh30 and SMS-CTR cells (**Figure 4a**). We also detected overlapping CTCF and SNAI2 binding peaks at the 5’ *PAX3* TAD boundary region (**Figure 4b**). We found that a subset of SNAI2 binding peaks in FN-RMS, SMS-CTR and RD cells, overlapped with CTCF at predicted TAD boundaries. We hypothesized that this CTCF/SNAI2 overlapping site was important in defining the TAD boundary and may also be important for maintaining *PAX3* expression in RMS cells. (**Figure 4b**). We next asked whether *PAX3* served essential roles in RMS and noted a differential requirement for expression of *PAX3* in DepMap (66) with it being essential in FP-RMS cells, while not required for viability in FN-RMS cells (**Figure 4c**). However, it is challenging to distinguish whether the DepMap guide RNAs are able to target endogenous *PAX3* versus the *PAX3-FOXO1* fusion gene. We have previously observed that *PAX3-FOXO1* is an essential gene in Rh4, while the loss of the fusion gene has lesser effects on Rh30 (67). Thus, to investigate the functional roles of chromatin domains for essentiality of *PAX3* expression, we designed CRISPR/Cas9 guides to delete the TAD boundary at the CTCF/SNAI2 site at in Rh30 cells. We used a two guide CRISPR/Cas9 strategy and deleted a 91bp region upstream of the *PAX3* gene that contained the 5’ TAD boundary in Rh30, and also in SMS-CTR cells. We confirmed the deletion by genomic PCR (**Figure 4d**) and performed western blot assays to assess PAX3 protein expression. We observed a >90% loss of PAX3 protein when the *PAX3* TAD-boundary was deleted in SMS-CTR and Rh30 cells, implying that this TAD boundary element is essential for endogenous *PAX3* expression. We next assessed expression of the PAX3-FOXO1 fusion protein using both the PAX3 and the FOXO1 antibodies. We observed that FOXO1 as well as PAX3-FOXO1 expression was significantly increased in Rh30 cells upon deletion of the boundary element (**Figure 4e**). This is consistent with previous studies showing mutual exclusivity in expression of PAX3-FOXO1 relative to PAX3 (68,69). Interestingly, in SMS-CTR cells the endogenous FOXO1 protein was also increased (**Figure 4e**).

**Figure 4.**
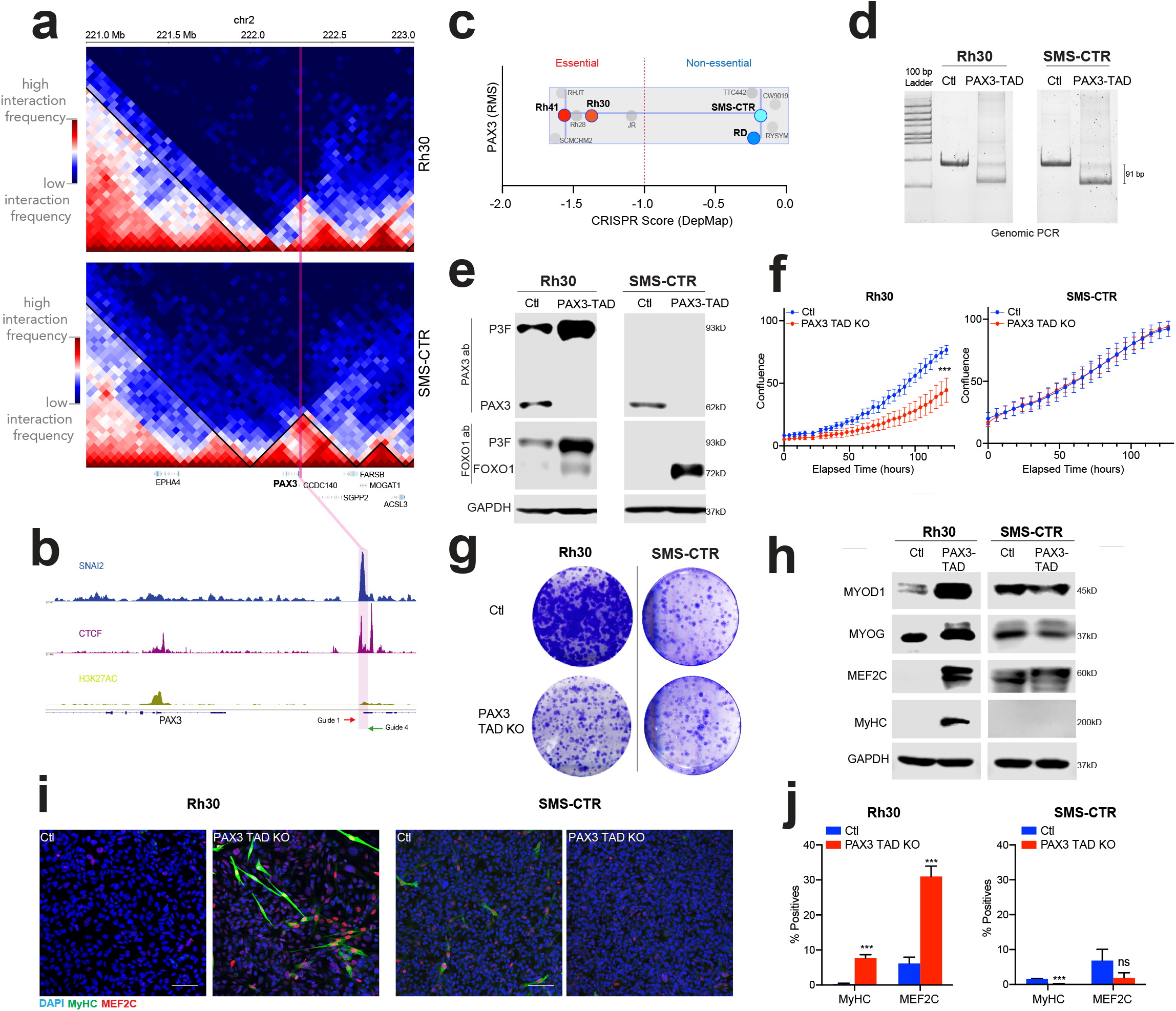
Functional requirements for TADs. (**a**) HiC data corresponding to the *PAX3* TAD region is shown, for SMS-CTR and Rh30 at Chromosome 2 (**b**) ChIP-seq for CTCF, SNAI2, and H3K27ac at the *PAX3* locus in SMS-CTR cells. CRISPR guide-RNAs targeting the TAD at this locus is indicated with *Guide 1-4*. (**c**) DepMap CRISPR score for RMS cell lines, values below -1 indicate essentiality of *PAX3*. (**d**) Genomic PCR of the CRISPR/Cas9 targeted region in control and *PAX3*-TAD deleted cells (**e**) Expression of PAX3, FOXO1, PAX3-FOXO1 (P3F) and GAPDH control in Rh30 and SMS-CTR cells, with SNAI2-CTCF peak at *PAX3* TAD boundary deleted (*PAX3*-TAD KO). (**f**) Cell growth analysis of *PAX3*-TAD deleted cells compared to control Rh30 and SMS-CTR cells. (**g**) Colony assay of *PAX3*-TAD deleted cells compared to control Rh30 and SMS-CTR cells. (**h**) Expression of myogenic differentiation factors in *PAX3* TAD deleted compared to control in Rh30 and SMS-CTR cells. (**i**) Confocal images of Control and *PAX3*-TAD deleted cells after three days of differentiation, cells were stained for DAPI (blue), differentiated myosin MyHC (green) and MEF2C (red). (**j**) Quantification of immunostaining from *PAX3* TAD deleted cells compared to control cells grown in differentiation media.

We next assessed the functional consequences of the loss of the *PAX3* TAD boundary. We performed growth assays, and found that while Rh30 cells are viable, they have a deficit in growth compared to control cells, while in SMS-CTR cell growth was unaffected (**Figure 4f**). In colony forming assays, we saw a similar trend, where the *PAX3* TAD boundary was essential for growth, while SMS-CTR cells were not affected **(Figure 4g**). Lastly, we assessed the effect of the loss of *PAX3* on myogenic differentiation. In Rh30 cells, loss of PAX3 expression, through deletion of its TAD boundary, resulted in an increase in MYOD, MYOG, MEF2C, and differentiated myosin MyHC expression (**Figure 4h**), while SMS-CTR cells did not show an induction of myogenic differentiation regulators. This effect was also observed in cells grown in myogenic differentiation media, where Rh30 *PAX3*-TAD deleted cells showed increased myogenic differentiation (**Figure 4i,j**), while SMS-CTR cells did not show any increase in myogenic differentiation. Together, we demonstrate that our identification of 3D chromatin structures at the *PAX3* locus reveals essential components of its transcriptional and oncogenic regulation in rhabdomyosarcoma. Furthermore, by selectively targeting the *PAX3* TAD we can selectively deplete *PAX3* and not *PAX3-FOXO1* expression thus illuminating roles of *PAX3* in the tumor.

## DISCUSSION

We report the genome structure of RMS in each major subtype. In our Hi-C studies, we observe common and distinct architectural and epigenetic events across FN and FP RMS subtypes. We anticipate further advances in this area as the epigenetics and genome structure associated with the infrequent *PAX3-NCOA*, and *PAX3-INO80* fusions in RMS are elucidated and compared with the *PAX3-FOXO1* and *PAX7-FOXO1* fusion drivers (4). In our studies we identify unique structural elements underlying large scale compartmental structures, TADs, and structural variations. In the cases of compartments, we find tumor-driving TFs highly associated with each class of compartment category (A, and B). In our analyses of TADs, we find a surprising lack of co-occurrence of MYOD and PAX3-FOXO1 binding sites flanking these structural elements, while our evidence supports that the CTCF “rule” of flanking convergent motifs applies strongly in each major subtype of the tumor. In the case of CNV, we find evidence from our Hi-C data that gene loci encoding *CDK4, MYCN*, and *PAX3-FOXO1* are amplified suggesting that multiple pathways may reinforce the genetics driving tumorigenesis (2,27).

It is interesting to note that SVs are not shared between cell lines, even within a subtype. We did not detect any subtype-specific SVs that are common to both FN-RMS cell lines. In FP-RMS cell lines, the only common SV we detected is the *PAX3-FOXO1* fusion locus. These results motivated the question, what chromatin domains are functionally driving in FP-RMS? To further evaluate this question, and the structural elements within the *PAX3-FOXO1* locus in Rh30, we performed CRISPR deletion with guide RNAs targeting CTCF and SNAI2-co-bound sites within the neo-TAD (**Figure 4a,b**). In these experiments, we found that PAX3 expression was highly dependent on the CTCF/SNAI2 sites at the TAD boundary (**Figure 4c,d**). The effects of PAX3 loss in these cells was minor decrease in proliferation and significant induction of myogenic differentiation in FP-RMS (**Figure 4f-h**). Thus, our chromatin structural studies have helped us define structure-function relationships in FP-RMS, within the *PAX3* - TAD. Further studies will be important to understand other “functional” TADs in RMS, within each major subtype, as previous studies have shown a decoupling of TAD structure and RNA Pol II activity (34).

Interestingly, we and others (29) have identified many SV events in the fusion negative subtype. Given the prevalence of structural variation in FN-RMS, and the new data indicating lack of clinical severity of *RAS* mutations in this subtype (2), we hypothesize that there may be fusion genes in FN-RMS cell lines. For example, our evidence suggests that there exists excessive interaction between *CGGBP1* on chr3 and *ATP8A2* on chr13 in SMS-CTR cell line (**Supplementary Figure S11**). Long-read RNA-sequencing and high content-imaging (26,70) will be efficacious in further studies of potential fusion driver events of embryonal RMS, to expand the scope of known alterations in this tumor. Moreover, future studies will be impactful for domain-specific normalizations in the context of spike-in chromatin (**Supplementary Figure S12**).

To the best of our knowledge, our study defines chromatin architecture in RMS at high depth for the first time. While there are no precision therapies yet reported for this tumor, more precise mechanistic and structural understanding of the chromatin initiation and maintenance in RMS will reveal new possibilities for precision therapy. It is of note that structural features we have identified are also functional features, reinforcing tumor growth at several key levels. Understanding the chromatin structural context for new driver genes, as they emerge, and transcriptional mechanisms along the linear chromatin fiber will be impactful in the RMS research community and within the larger community of oncologists and molecular biologists seeking to understand RMS etiology.

## DATA AVAILABILITY

The data discussed in this publication have been deposited in NCBI’s Gene Expression Omnibus and are accessible through GEO Series accession number GSE215203 (https://www.ncbi.nlm.nih.gov/geo/query/acc.cgi?acc=GSE215203). Previously published SRA datasets used in the study were: ChIP-seq and RNA-seq in RMS (GSE83728). A source data file accompanies this manuscript. The remaining data are available within the article, supplementary Information or available from the authors upon request.

## SUPPLEMENTARY DATA

### Spice-C Protocol (Spiked-in Chromatin Equalized HiC)

RMS cells grown (10 and 12 million cells) in tissue culture dishes were treated with Trypsin and resuspended in fresh growth media. For 13 mL of counted cells in culture medium, add 360 μL of 37% (wt/vol) formaldehyde. Quench the formaldehyde with a final concentration of 125 mM glycine at 4 °C for 5 min. Pellet cells at 1,250g for 3 min at 4 °C. Remove and discard the supernatant. Resuspend in cold PBS (5 mL) on ice, pellet again at 1,250g for 3 min at 4 °C and discard the supernatant. Prepare exogenous spiked-in cells from an orthogonal species in the same manner as described and store at - 80 °C or proceed to lysis.

### Lysis and restriction digest

Resuspend 6 million crosslinked human cells mixed with 2 million crosslinked mouse cells (∼20% orthologous chromatin, by cell equivalency for SPICE-C) in 500 μL of ice-cold Hi-C Lysis Buffer (with freshly added protease inhibitor cocktail) and rotate at 4 °C for 30 min. Centrifuge at 2,500g for 5 min at 4 °C, discard the supernatant. Wash pellet with 500 μL of ice-cold Hi-C Lysis Buffer (without resuspending it). Centrifuge again at 2,500g for 5 min at 4 °C. Remove and discard the supernatant and resuspend in 100 μL of 0.5% (wt/vol) SDS (in TE pH 7.4) to permeabilize nuclei in preparation for in situ enzymatic digestion steps. Incubate mixture for 10 min at 62 °C and then add 285 μL of water and 50 μL of 10% (vol/vol) Triton X-100 to quench the SDS. Mix by pipetting, spin down the sample and incubate for 15 min at 37 °C. Add 50 μL of 10× NEBuffer 2, mix well and then add 200 U (8 μL) of MboI restriction enzyme, mix and digest chromatin for 2 h at 37 C with rotation or agitation. Heat inactivate MboI for 20 min at 62 °C. Then, cool the sample at 4 °C for 5 min.

### Biotin incorporation and proximity ligation

After in situ enzymatic digestion with MboI, overhangs will be blunted and biotinylated. To the heat-inactivated mixture from, add 52 μl of the following master mix: 37.5 μL 0.4 mM biotin-dATP, 1.5 μL 10 mM dCTP, 1.5 μL 10 mM dGTP, 1.5 μL 10 mM dTTP, 10 μL 5U/μL DNA Polymerase I, Large (Klenow) Fragment (NEB, M0210). Mix gently by pipetting and incubate at 37 °C for 1 h with gentle agitation in a ThermoMixer. Prepare the ligation mix (T4 ligase, 10x buffer) and add to samples mix well and proceed to the next step. Incubate at room temperature for 4 h with gentle agitation in a ThermoMixer (300 r.p.m). Centrifuge reaction mixture at 2,500g for 5 min at 4 °C and remove and discard the supernatant. Use pelleted nuclei in subsequent steps.

### Sonication

Bring pellet up to 700 μL in TE supplemented with freshly added protease inhibitor cocktail. Use 10 min of total shearing ‘on’ time with 30 s ‘on’ and 30 s ‘rest’ with an Active Motif Epi-shear probe sonicator (see Equipment) at 30% power. While checking the shearing efficiency, keep samples at 4 °C. Add 15 μL of 10% SDS for 600 μL of lysate (2.5 μL 10% SDS/100 μL) mix by pipetting and add 10 μL of Proteinase K, per tube and incubate at 65 °C O/N with shaking in a benchtop ThermoMixer at 500– 700 r.p.m. Purify the DNA from the samples with two Minelute columns and elute in 20 μL per column with elution buffer provided in the kit. Combine the eluates to a total of ∼ 36 μL and quantify DNA, use 5-10 μg diluted in TE for Biotin capture.

### Biotin capture

Prepare for biotin pulldown by washing 50 μL of M-280 Streptavidin Dynabeads (C) twice with 50 μL of Tween Wash Buffer. Resuspend the beads in 50 μL of 2X Biotin Binding Buffer and add to the 50 μL sample diluted in TE. Incubate the mixture at room temperature for 30 min with rotation. Let stand on the magnet and remove and discard the buffer. Resuspend the beads in 500 μL of Tween Wash Buffer and incubate at 55 °C for 2 min with shaking at 300–400 r.p.m.; then, wash with 500 μL of TE pH 7.4 and leave at room temperature for on-bead library preparation.

## ACKNOWLEDGEMENT

We would like to thank Amy Wetzel, Huachun Zhong, Shireen Woodiga, and the Institute for Genomic Medicine sequencing facility at Nationwide Children’s Hospital for sequencing expertise and guidance. We would like to thank Dr. Peter Houghton for deriving the rhabdomyosarcoma cell lines used in this study. Dr. Åsa Karlström, and Childhood Solid Tumor Network at St. Jude Children’s Research Hospital provided the rhabdomyosarcoma PDX samples, and Dr. Ruoning Wang provided mouse muscle tissue for methods development. We wish to thank our colleagues at Nationwide Children’s Hospital and the OSU epigenetics community, UT Health, San Antonio, and NCI-CCR for conversations that were helpful in developing this work.

## TABLE AND FIGURES LEGENDS

**Supplementary Figure S1. Extended data: Rh4 compartments**. The associations of GC-richness, H3K27ac, H3K9me3, MYOD binding, and PAX3-FOXO1 binding are shown for Rh4 cells on all chromosomes with A-compartments shown in blue and B-compartment shown in black.

**Supplementary Figure S2. Extended data: Rh30 compartments**. The associations of GC-richness, H3K27ac, H3K9me3, MYOD binding, and PAX3-FOXO1 binding are shown for Rh30 cells on all chromosomes with A-compartments shown in blue and B-compartment shown in black.

**Supplementary Figure S3. Extended data: RD compartments**. The associations of GC-richness, H3K27ac, H3K9me3, and MYOD binding are shown for RD cells on all chromosomes with A-compartments shown in blue and B-compartment shown in black.

**Supplementary Figure S4. Extended data: SMS-CTR compartments**. The associations of GC-richness, H3K27ac, H3K9me3, and MYOD binding are shown for SMS-CTR cells on all chromosomes with A-compartments shown in blue and B-compartment shown in black.

**Supplementary Figure S5**. Saddle Plots showing A-A, A-B and B-B compartmental interaction strength in (a) Rh4, (b) Rh30, (c) RD and (d) SMS-CTR cell lines.

**Supplementary Figure S6. Correlation between PC1 values and chromatin factor binding sites**. PC1 values for compartments are plotted against *number of binding sites* for chromatin marks and regulatory factors, H3K27ac as scatter (**a**) and violin plots (**e**), H3K9me3 as scatter (**b**) and violin plots (**f**), MYOD as scatter (**c**) and violin plots (**g**), and PAX3-FOXO1 as scatter (**d**) and violin plots (**h**).

**Supplementary Figure S7. Extended data: subtype-specific compartments in RMS**. Compartment calls contrasting A and B compartments for each chromosome (not in order) respectively for Rh4, Rh30, RD, and SMS-CTR cell lines.

**Supplementary Figure S8. Differential TAD structure at gene:** (**a**) *TCF4*, (**b**) *PTN*, and (**c**) *NRG1*.

**Supplementary Figure S9. Extended data: copy number from Hi-C**. Copy number variation (CNV) is indicated with colorimetric scale for Rh4, Rh30, RD, SMS-CTR cell lines for all chromosomes. On the scale, black = deletion/loss, blue = single copy, beige = two copies, and red = 3 copies. Fusion status is shown for each cell line, underneath the CNV plots.

**Supplementary Figure S10**. Interchromsomal and long-range intrachromosomal interactions at PAX3-FOXO1 break site. We observed interchromsomal interaction between chr2:219Mb-222.5Mb and chr13:77Mb, as well as long-range intrachromosomal interaction between chr13:40.6Mb and chr13:77Mb.

**Supplementary Figure S11**. Evidence of fusion between genes *CGGBP1* on chr3 and *ATPBA2* on chr13 in SMS-CTR cell lines.

**Supplementary Figure S12. Hi-C library preparations**. 2% agarose E-Gel EX images are shown for RD, SMS-CTR (CTR), Rh30, and Rh4 cells through the steps of Hi-C library generation. In each case, the DNA ladder at left is marked from bottom to top with 100 bp, 200 bp, 300 bp, 400 bp, 500 bp, 600 bp, 700 bp, 800, bp, 900 bp, 1kb, 1.5 kb, 2kb, 2.5 kb, 3kb, 4kb, 5kb, 6kb, 8kb, 10kb. (**a**) Lysis and sonication are shown during the *in situ* steps. (**b**) Biotinylation and proximity ligation are represented for each cell line. (**c**) PCR with condition-specific indexes is shown for each cell line, indicating the fragment length distribution after purification. (**d**) Library preparation steps including lysis, MboI digest, biotinylation, proximity ligation, and sonication are shown for PDX sample SJRHB011. (**e**) Library amplification with indexed primers is shown for PDX sample SJRHB011. (**f**) Biotin capture optimization is shown for 4 ug and 8 ug DNA, respectively, for RD cell Hi-C library preparation. (**g**) Library preparation steps including lysis, MboI digest, biotinylation, proximity ligation, and sonication are shown for PDX sample SJRHB013758 (left) and SJRHB013 (right). Library amplification with indexed primers is shown for PDX samples SJRHB013758 (left) and SJRHB013 (right) before (**h**) and after (**i**) purification.

**Supplementary Table S1**. Myogenic and neurogenic genes (from GSEA) associated with discordant compartments.

**Supplementary Table S2**. Neurogenic genes in differential TADs.

**Supplementary Table S3**. CNV calls for Rh4, Rh30, RD, SMS-CTR cell lines annotated to the closest genes.

