## Supplemental Figures for "The 3D chromatin landscape of rhabdomyosarcoma": Figure S6.pdf

### Figure S6. Correlation of PC1 values with chromatin factor binding sites (Rh4 Cell Line).

Hi-C PC1 value association with binding sites of histone marks and tumor-essential factors

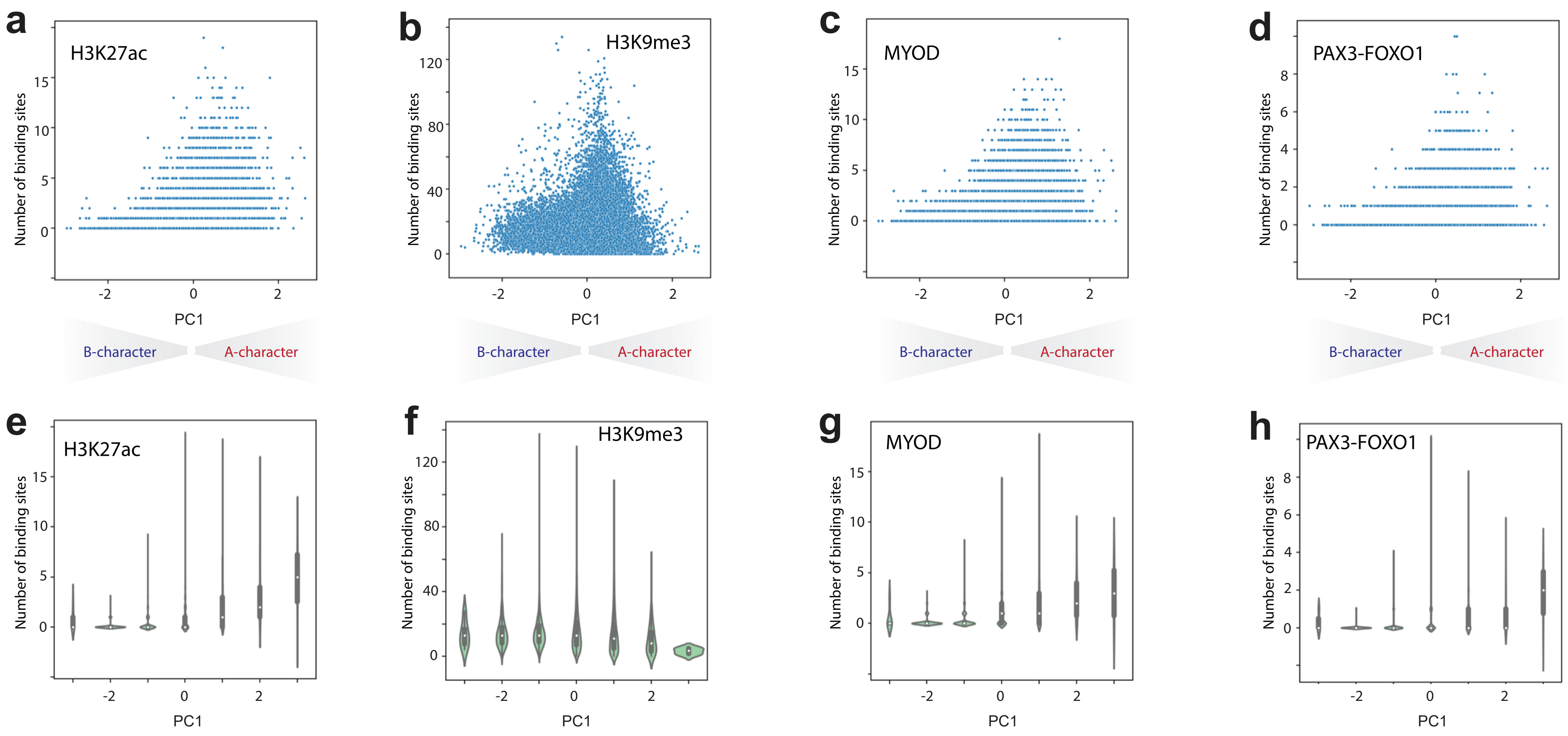
