## Supplementary figures and images for "The 3D chromatin landscape of rhabdomyosarcoma"

### Figure S1.pdf

#### Figure S1. Extended data: Rh4 compartments.

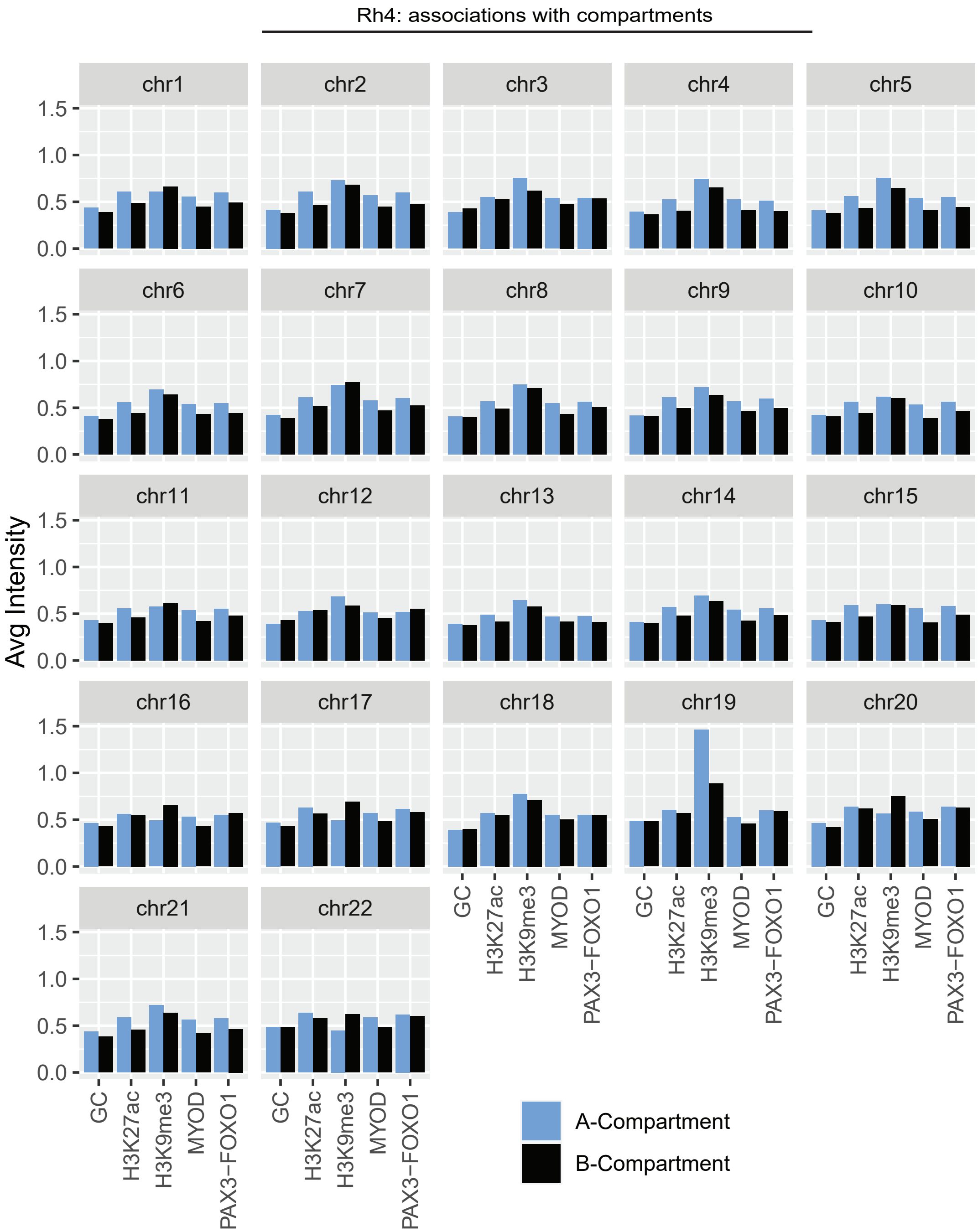

### Figure S2.pdf

#### Figure S2. Extended data: Rh30 compartments.

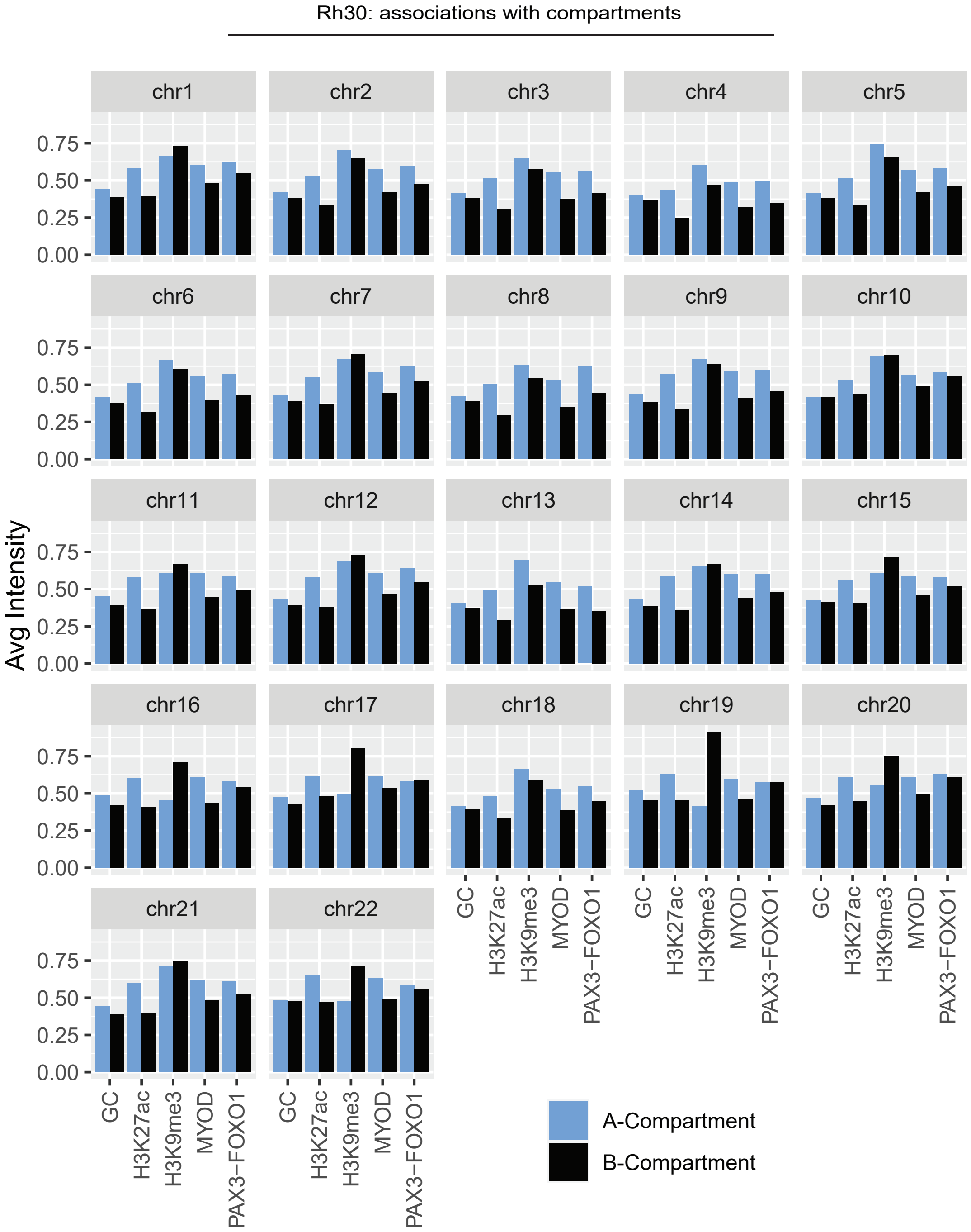

### Figure S3.pdf

#### Figure S3. Extended data: RD compartments.

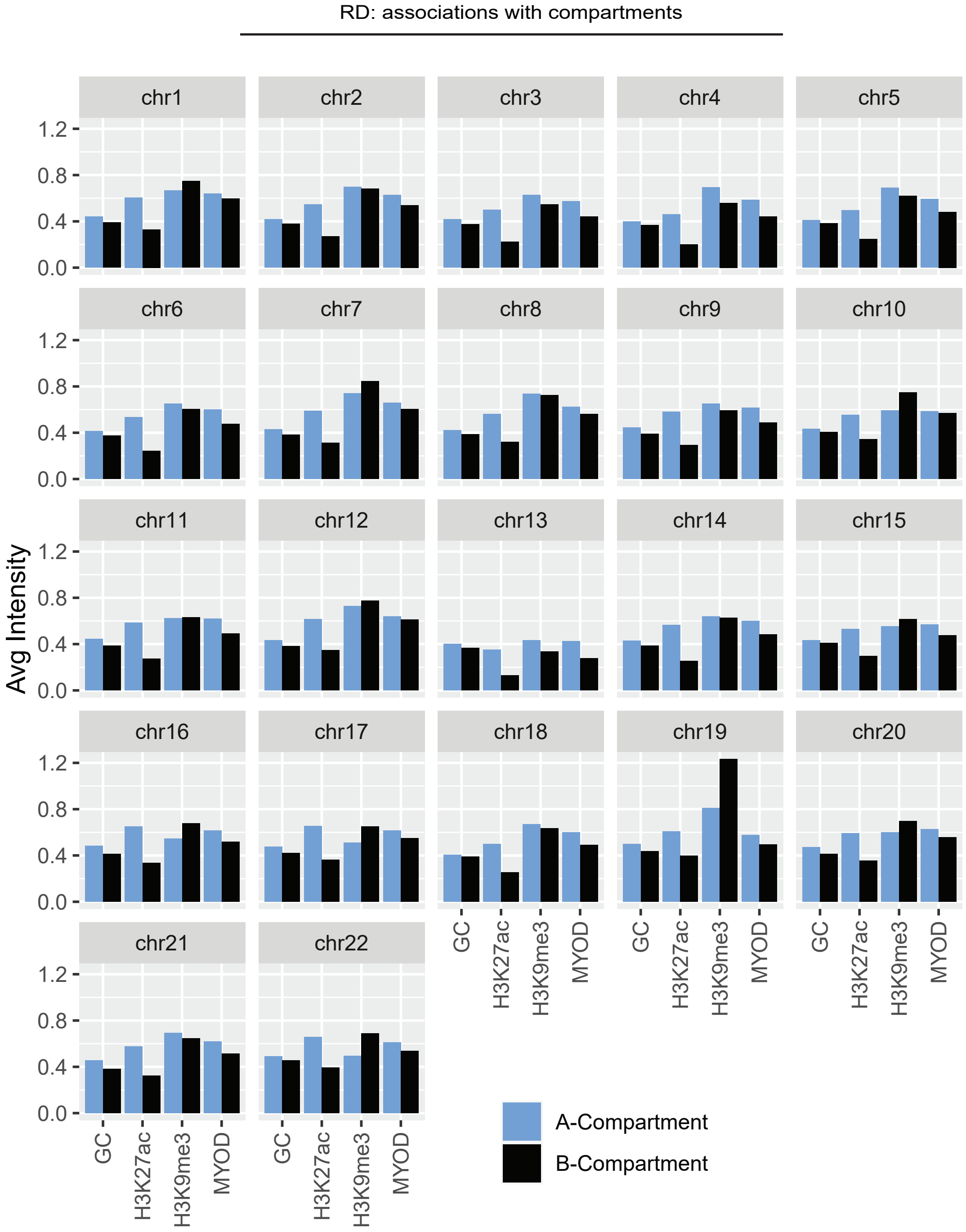

### Figure S4.pdf

#### Figure S4. Extended data: SMS-CTR compartments.

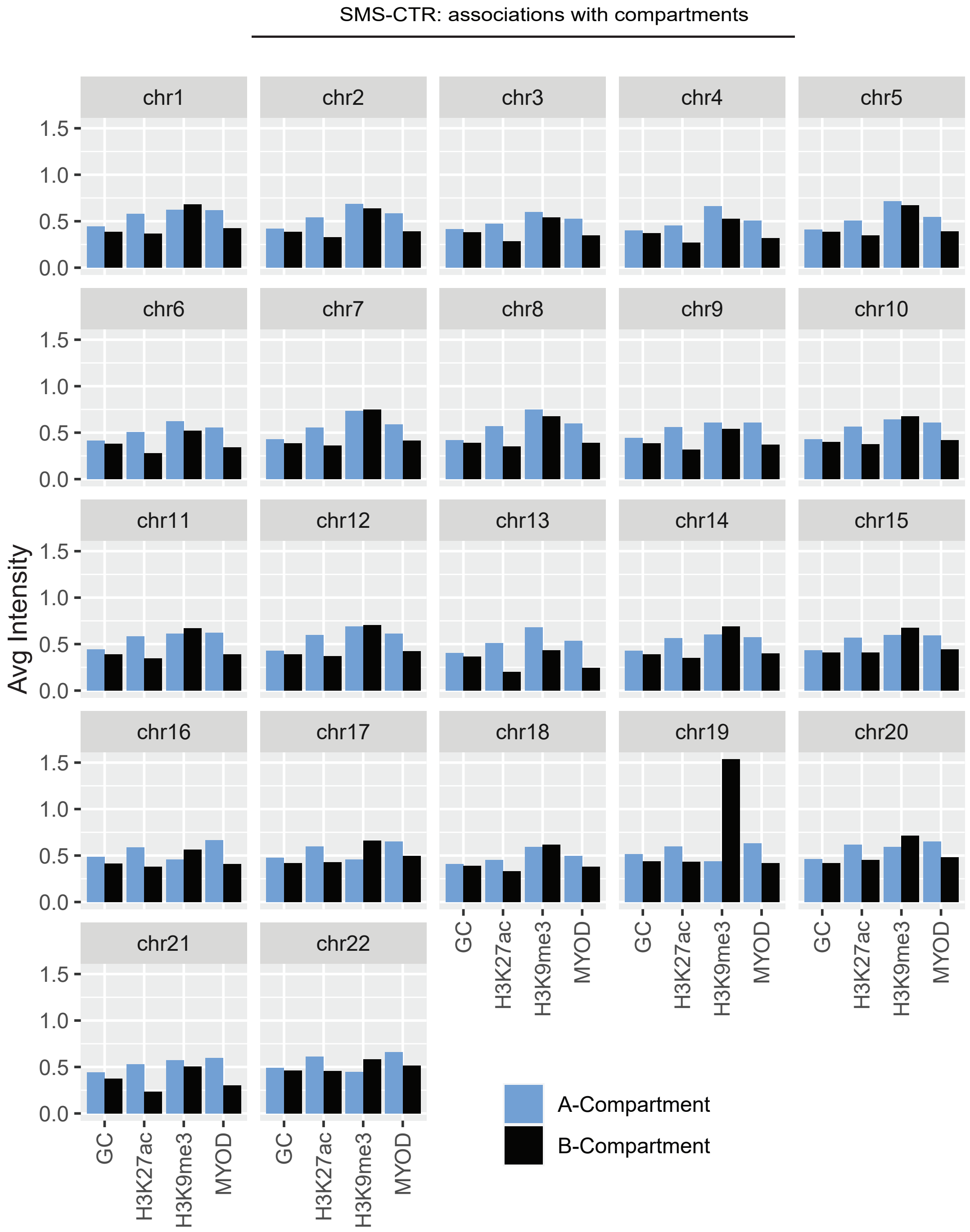

### Figure S5.pdf

#### Figure S5. Extended data: Saddle Plots.

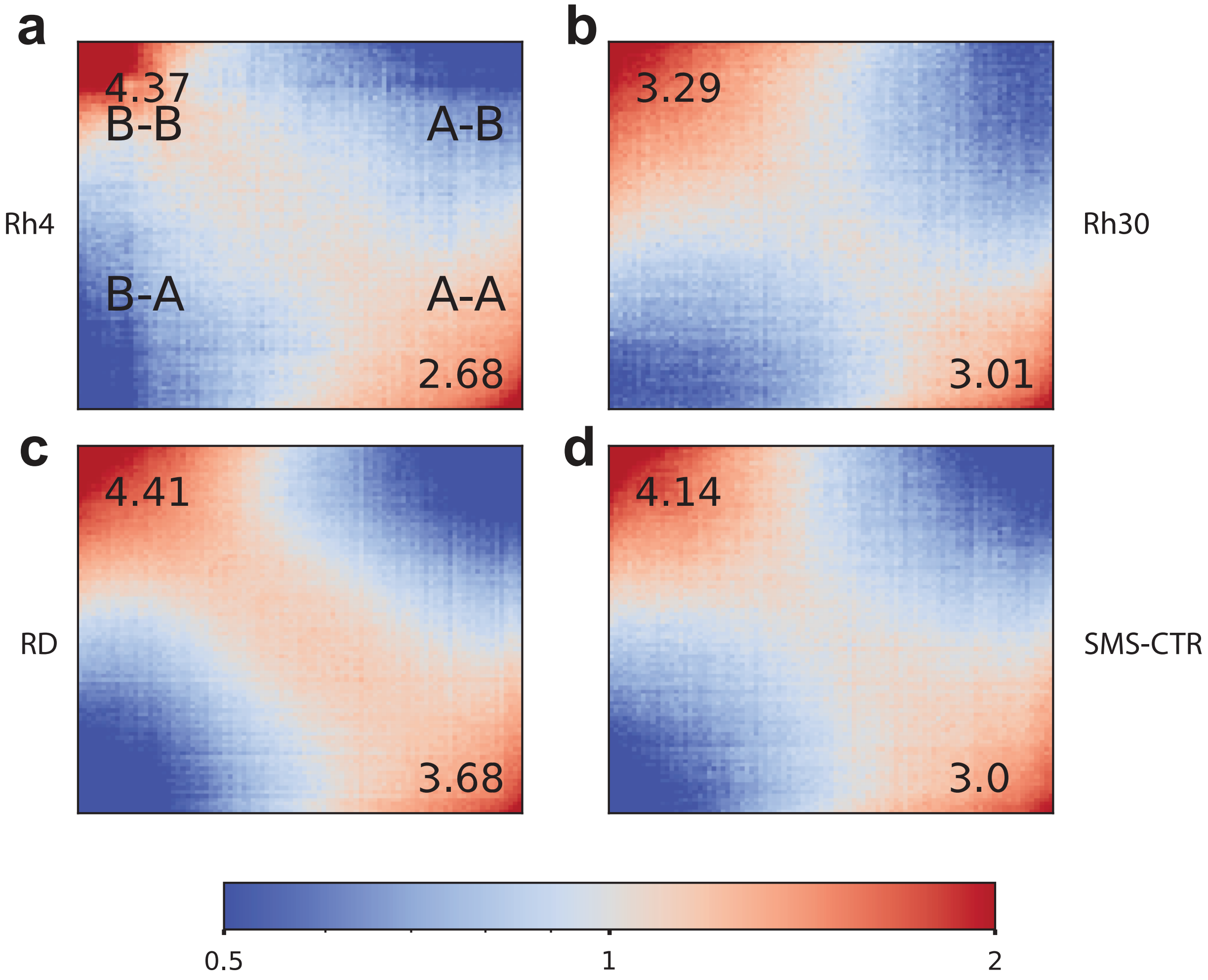

### Figure S7.pdf

#### Figure S7. Extended data: subtype-specific compartments in RMS.

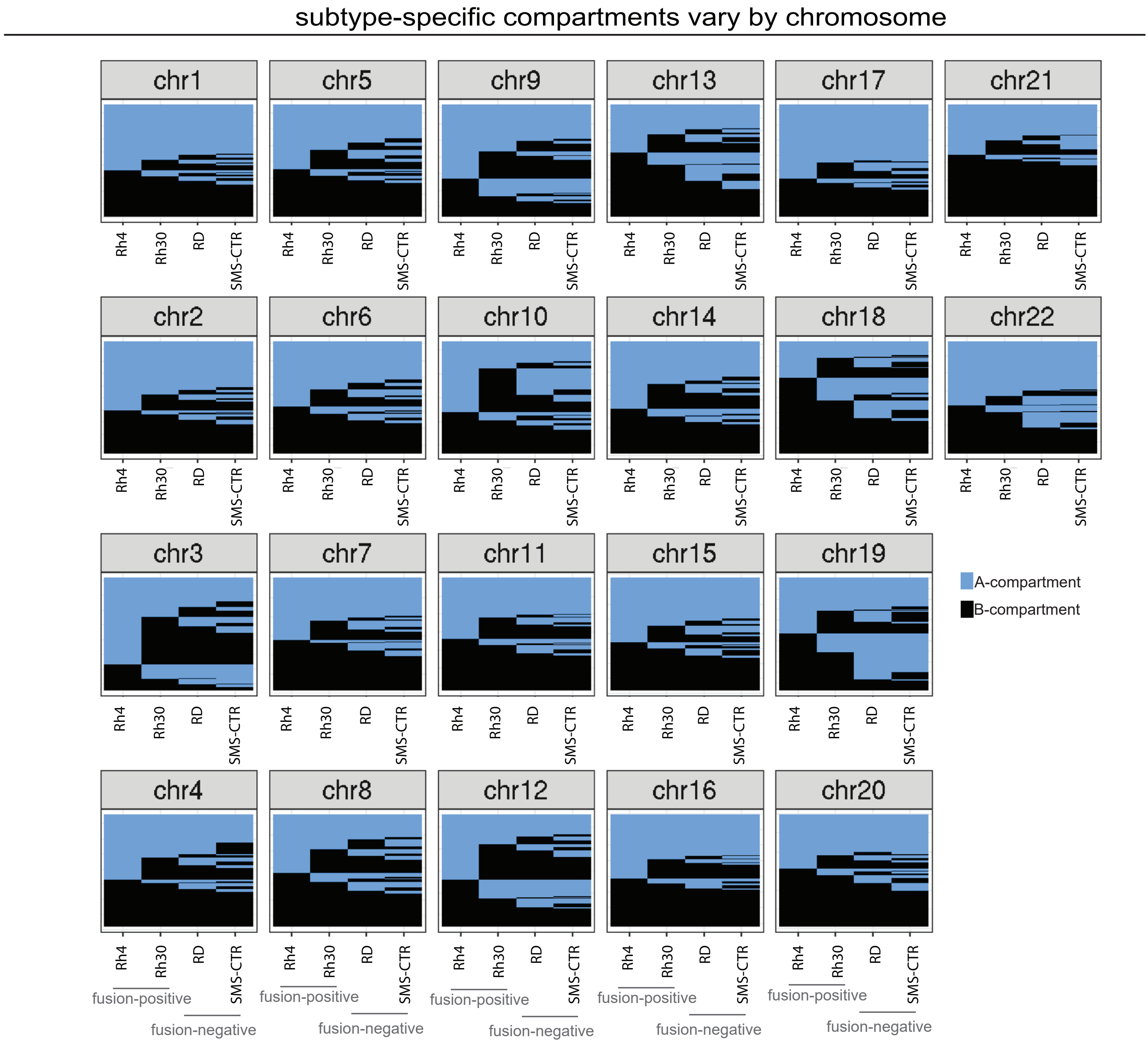

### Figure S8.pdf

Figure S8. Extended data: Differential TADs

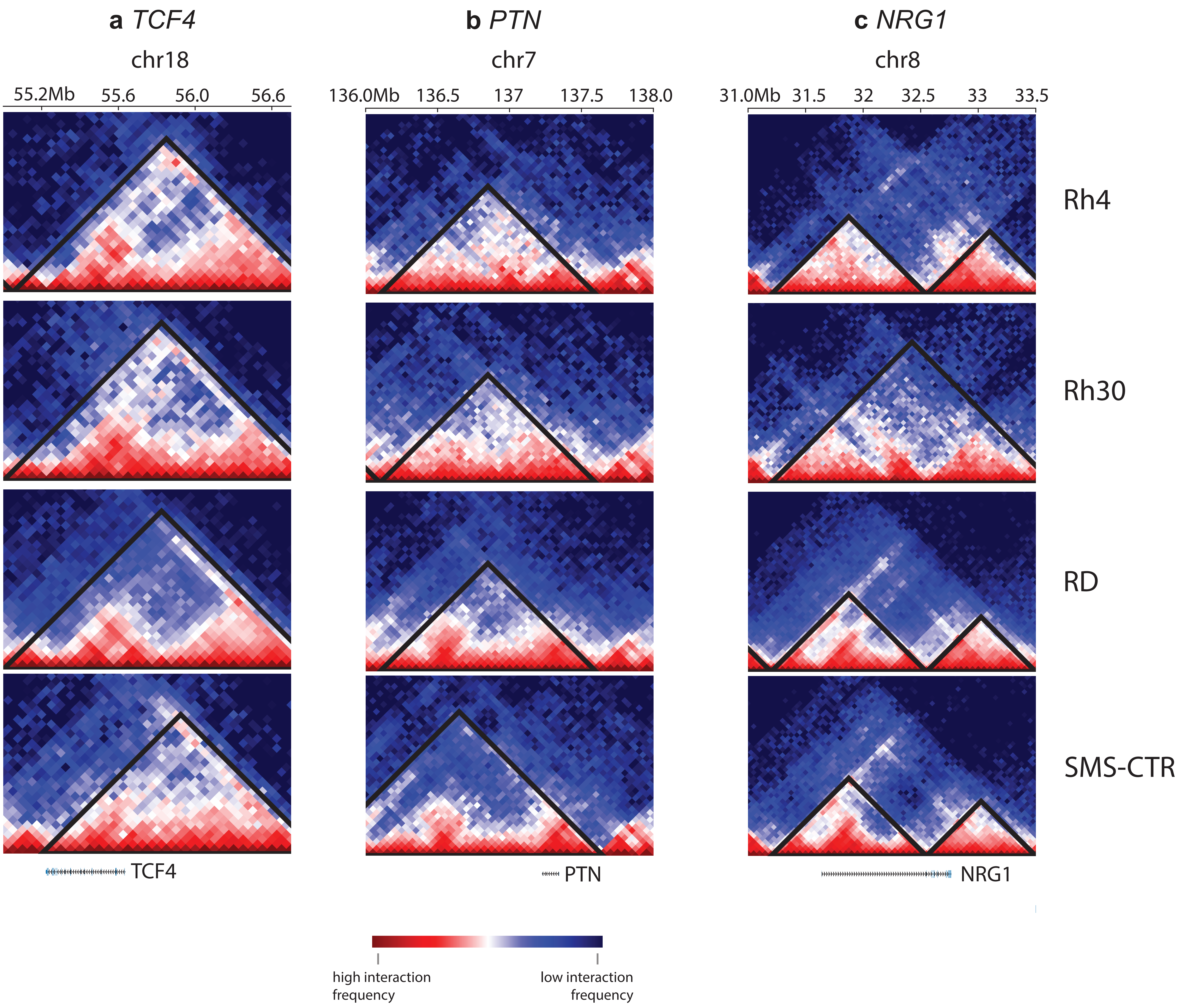

### Figure S9.pdf

#### Figure S9. Extended data: copy number from Hi-C.

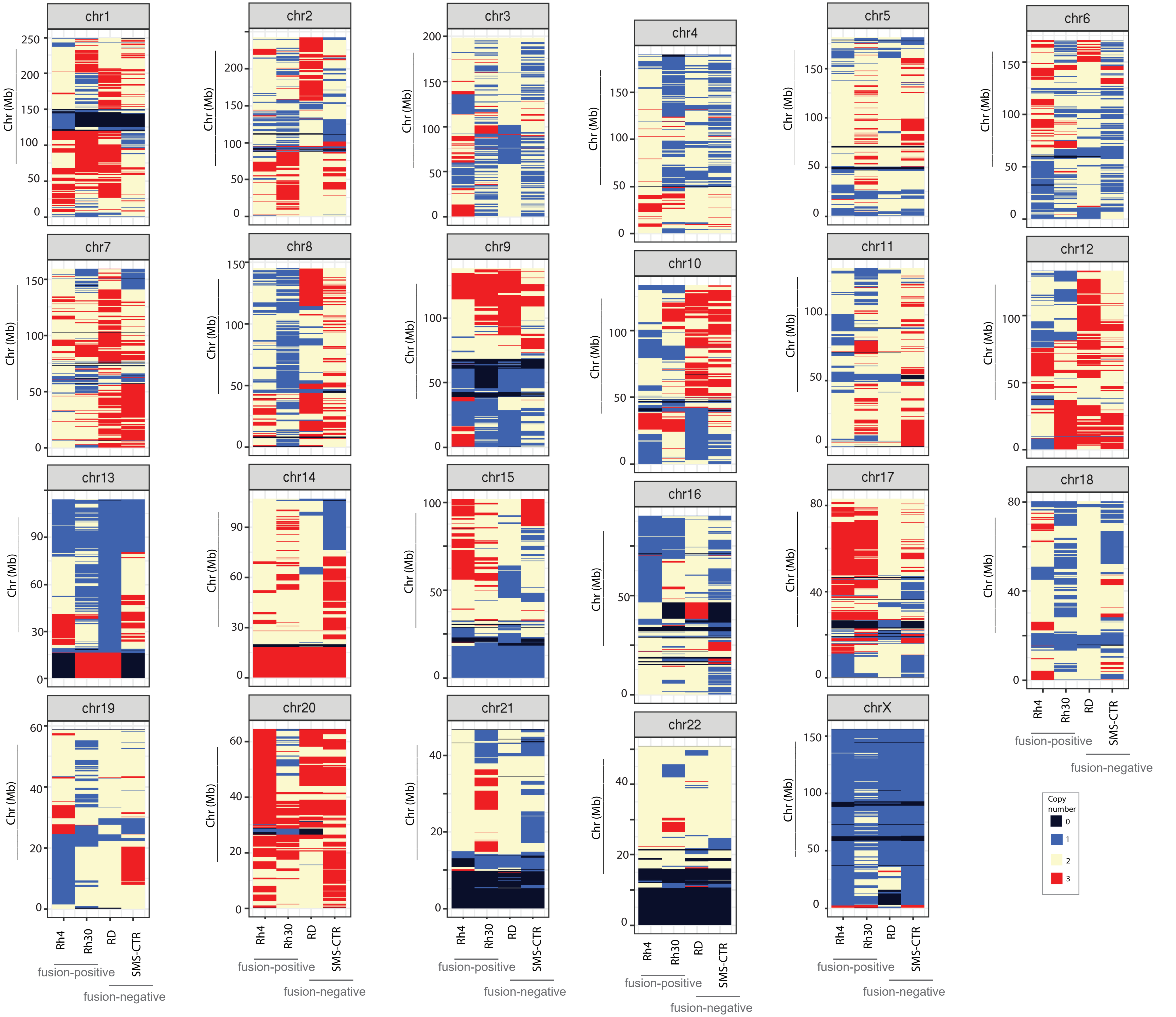

### Figure S10.pdf

#### Figure S10. Extended data: long-range/interchromosomal interaction in Rh4

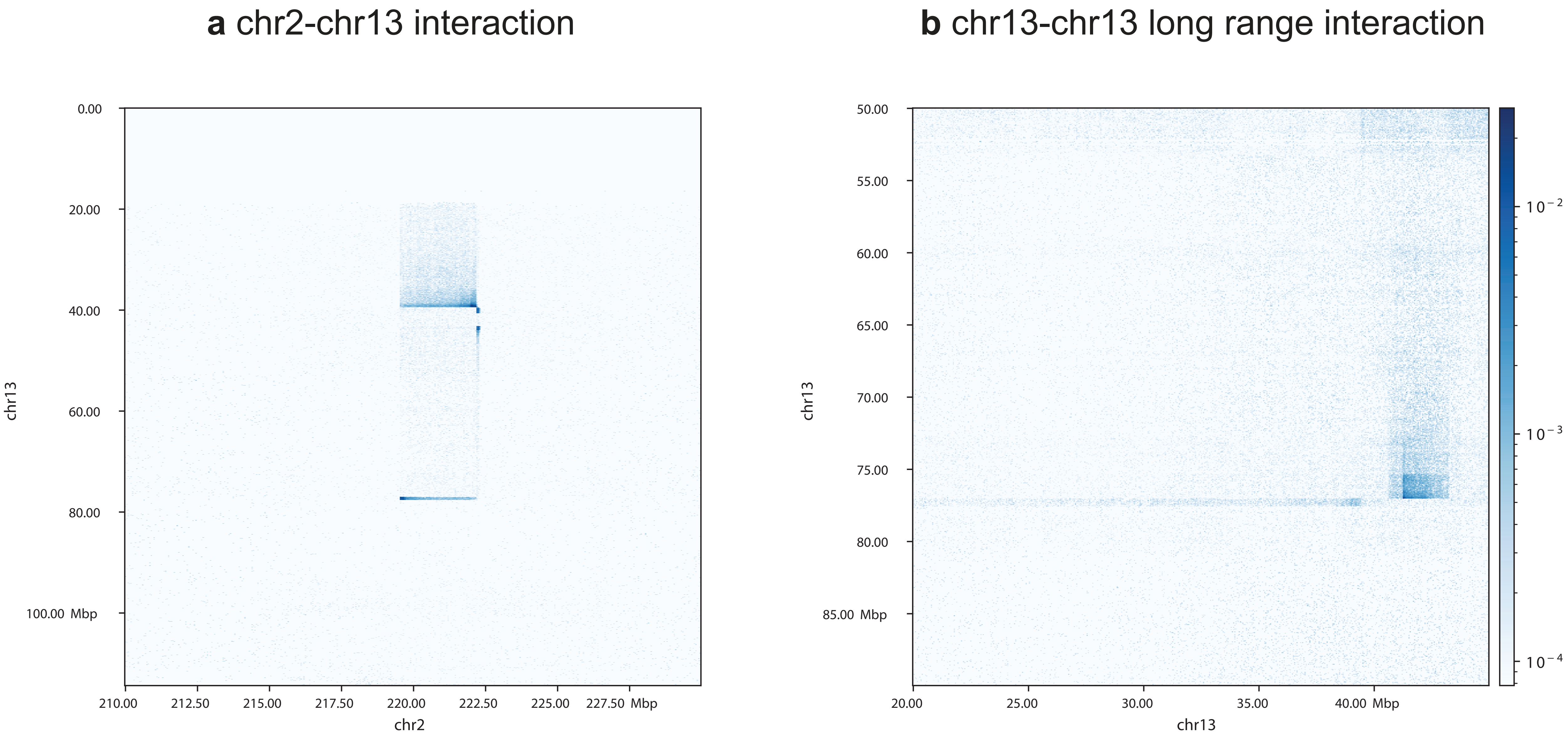

### Figure S11.pdf

#### Figure S11. Extended data:
#### Evidence of fusion between chr3 and chr13 (SMS-CTR)

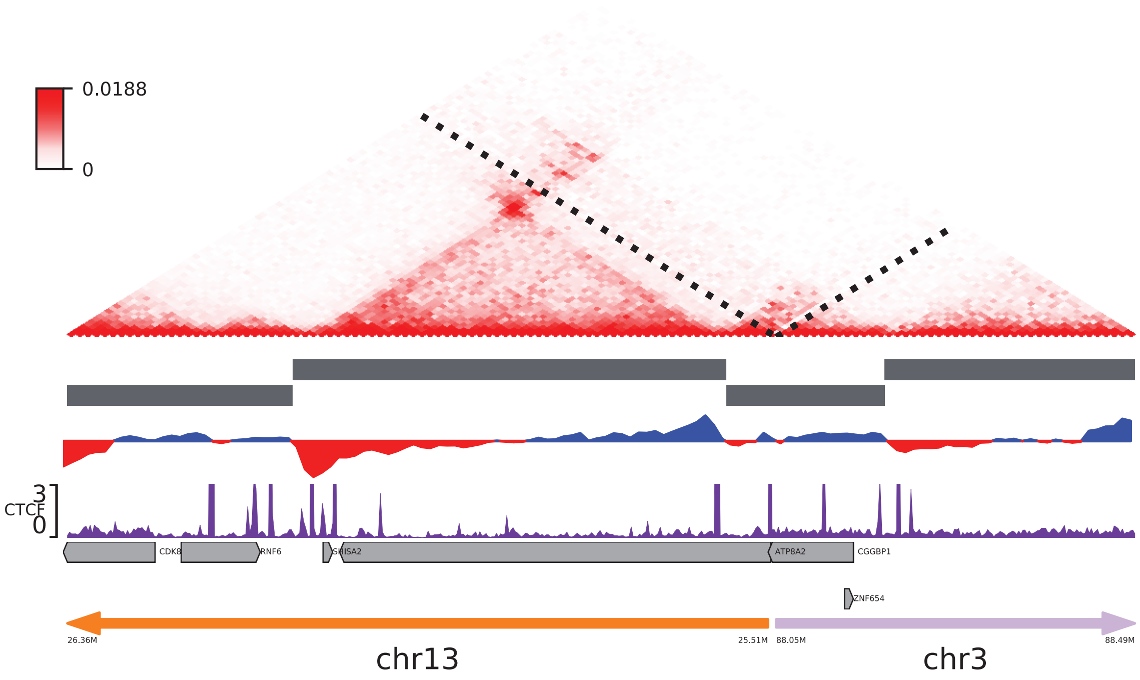

### Figure S12.pdf

#### Figure S12. Hi-C library preparations.

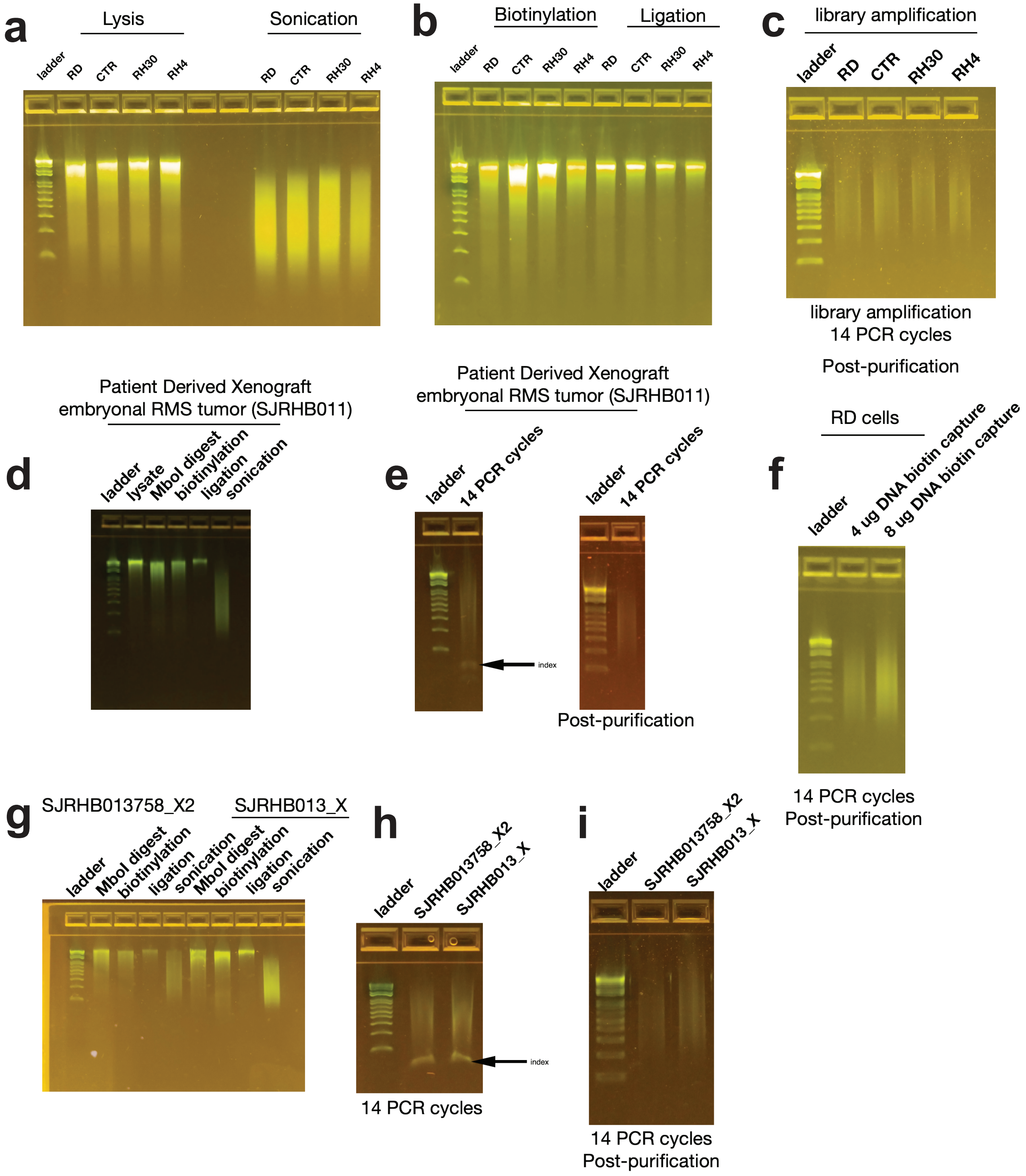
